# FRETBursts: An Open Source Toolkit for Analysis of Freely-Diffusing Single-Molecule FRET

**DOI:** 10.1101/039198

**Authors:** Antonino Ingargiola, Eitan Lerner, SangYoon Chung, Shimon Weiss, Xavier Michalet

## Abstract

Single-molecule Förster Resonance Energy Transfer (smFRET) allows probing intermolecular interactions and conformational changes in biomacromolecules, and represents an invaluable tool for studying cellular processes at the molecular scale. smFRET experiments can detect the distance between two fluorescent labels (donor and acceptor) in the 3–10 nm range. In the commonly employed confocal geometry, molecules are free to diffuse in solution. When a molecule traverses the excitation volume, it emits a burst of photons, which can be detected by single-photon avalanche diode (SPAD) detectors. The intensities of donor and acceptor fluorescence can then be related to the distance between the two fluorophores.

While recent years have seen a growing number of contributions proposing improvements or new techniques in smFRET data analysis, rarely have those publications been accompanied by so.ware implementation. In particular, despite the widespread application of smFRET, no complete so.ware package for smFRET burst analysis is freely available to date.

In this paper, we introduce FRETBursts, an open source software for analysis of freely-diffusing smFRET data. FRETBursts allows executing all the fundamental steps of smFRET bursts analysis using state-of-the-art as well as novel techniques, while providing an open, robust and well-documented implementation. Therefore, FRETBursts represents an ideal platform for comparison and development of new methods in burst analysis.

We employ modern software engineering principles in order to minimize bugs and facilitate long-term maintainability. Furthermore, we place a strong focus on reproducibility by relying on Jupyter notebooks for FRETBursts execution. Notebooks are executable documents capturing all the steps of the analysis (including data files, input parameters, and results) and can be easily shared to replicate complete smFRET analyzes. Notebooks allow beginners to execute complex workflows and advanced users to customize the analysis for their own needs. By bundling analysis description, code and results in a single document, FRETBursts allows to seamless share analysis workflows and results, encourages reproducibility and facilitates collaboration among researchers in the single-molecule community.

## 1 Introduction

### 1.1 Open Science and Reproducibility

Over the past 20 years, single molecule FRET (smFRET) has grown into one of the most useful techniques in singlemolecule spectroscopy [1, 2]. While it is possible to extract information on sub-populations using ensemble measurements (e.g. [3,4]), smFRET unique feature is its ability to very straightforwardly resolve conformational changes of biomolecules or measure binding-unbinding kinetics in heterogeneous samples [5–9]. smFRET measurements on freely diffusing molecules (the focus of this paper) have the additional advantage, over measurements performed on immobilized molecules, of allowing to probe molecules and processes without perturbation from surface immobilization or additional functionalization needed for surface attachment [10,11].

The increasing amount of work using freely-diffusing smFRET has motivated a growing number of theoretical contributions to the specific topic of data analysis [12–24]. Despite this profusion of publications, most research groups still rely on their own implementation of a limited number of methods, with very little collaboration or code sharing. To clarify this statement, let us point that our own group’s past smFRET papers merely mention the use of custom-made software without additional details [16, 17]. Even though some of these software tools are made available upon request, or sometimes shared publicly on websites, it remains hard to reproduce and validate results from different groups, let alone build upon them. Additionally, as new methods are proposed in literature, it is generally difficult to quantify their performance compared to other methods. An independent quantitative assessment would require a complete reimplementation, an effort few groups can afford. As a result, potentially useful analysis improvements are either rarely or slowly adopted by the community. In contrast with other established traditions such as sharing protocols and samples, in the domain of scientific software, we have relegated ourselves to islands of non-communication.

From a more general standpoint, the non-availability of the code used to produce scientific results, hinders reproducibility, makes it impossible to review and validate the software’s correctness and prevents improvements and extensions by other scientists. This situation, common in many disciplines, represents a real impediment to the scientific progress. Since the pioneering work of the Donoho group in the 90’s [25], it has become evident that developing and maintaining open source scientific software for reproducible research is a critical requirement of the modern scientific enterprise [26,27]

Other disciplines have started tackling this issue [28], and even in the single-molecule field a few recent publications have provided software for analysis of surface-immobilized experiments [29–33]. For freely-diffusing smFRET experiments, although it is common to find mention of “code available from the authors upon reques” in publications, there is a dearth of such open source code, with, to our knowledge, the notable exception of a single example [34]. To address this issue, we have developed FRETBursts, an open source Python software for analysis of freely-diffusing single-molecule FRET measurements. FRETBursts can be used, inspected and modified by anyone interested in using state-of-the art smFRET analysis methods or implementing modifications or completely new techniques. FRETBursts therefore represents an ideal platform for quantitative comparison of different methods for smFRET burst analysis. Technically, a strong emphasis has been given to the reproducibility of complete analysis workflows. FRETBursts uses Jupyter Notebooks [35], an interactive and executable document containing textual narrative, input parameters, code, and computational results (tables, plots, etc.). A notebook thus captures the various analysis steps in a document which is easy to share and execute. To minimize the possibility of bugs being introduced inadvertently [36], we employ modern software engineering techniques such as unit testing and continuous integration [28,37]. FRETBursts is hosted on GitHub [38, 39], where users can write comments, report issues or contribute code. In a related effort, we recently introduced Photon-HDF5 [40], an open file format for timestamp-based single-molecule fluorescence experiments. An other related open source tool is PyBroMo [41], a freely-diffusing smFRET simulator which produces Photon-HDF5 files that are directly analyzable with FRETBursts. Together with all the aforementioned tools, FRETBursts contributes to the growing ecosystem of open tools for reproducible science in the single-molecule field.

### 1.2 Paper Overview

This paper is written as an introduction to smFRET burst analysis and its implementation in FRETBursts. The aim is illustrating the specificities and trade-offs involved in various approaches with sufficient details to enable readers to customize the analysis for their own needs.

After a brief overview of FRETBursts features (section 2), we introduce essential concepts and terminology for smFRET burst analysis (section 3). In section 4, we illustrate the steps involved in smFRET burst analysis: (i) data loading (section 4.1), (ii) definition of the excitation alternation periods (section 4.2), (iii) background correction (section 4.3), (iv) burst search (section 4.4), (v) burst selection (section 4.6) and (vi) FRET histogram fitting (section 4.7). We conclude the section by surveying different methods proposed in littera-ture to study FRET dynamics (section 4.8). As an example of implementation of an advanced data processing technique, section 5 walks the reader through implementing Burst Variance Analysis (BVA) [23]. Finally, section 6 summarizes what we believe to be the strengths of FRETBursts software.

Throughout this paper, links to relevant sections of documentation and other web resources are displayed as “(link)”. In order to make the text more legible, we have concentrated Python-specific details in paragraphs titled *Python details.* These subsections provide deeper insights for readers already familiar with Python and can be initially skipped by readers who are not. Finally, note that all commands and figures in this paper can be regenerated using the accompanying notebooks (link).

## 2 FRETBursts Overview

### 2.1 Technical Features

FRETBursts can analyze smFRET measurements from one or multiple excitation spots [42]. The supported excitation schemes include single laser, alternating laser excitation (ALEX) with either CW lasers (μs-ALEX [43]) or pulsed lasers (ns-ALEX [44] or pulsed-interleaved excitation (PIE) [45]).

The software implements both standard and novel algorithms for smFRET data analysis including background estimation as a function of time (including background accuracy metrics), sliding-window burst search [10], dual-channel burst search (DCBS) [17] and modular burst selection methods based on user-defined criteria (including a large set of pre-defined selection rules). Novel features include burst size selection with Υ-corrected burst sizes, burst weighting, burst search with background-dependent threshold (in order to guarantee a minimal signal-to-background ratio [46]). Moreover, FRETBursts provides a large set of fitting options to characterize FRET subpopulations. In particular, distributions of burst quantities (such as *E* or *S*) can be assessed through (1) histogram fitting (with arbitrary model functions), (2) non-parametric weighted kernel density estimation (KDE), (3) weighted expectation-maximization (EM), (4) maximum likelihood fitting using Gaussian models or Pois-son statistic. Finally FRETBursts includes a large number of predefined and customizable plot functions which (thanks to the *matplotlib* graphic library [47]) produce publication quality plots in a wide range of formats.

Additionally, implementations of population dynamics analysis such as Burst Variance Analysis (BVA) [23] and two-channel kernel density distribution estimator (2CDE) [24] are available as FRETBursts notebooks (BVA link, 2CDE link).

### 2.2 Software Availability

FRETBursts is hosted and openly developed on GitHub. FRETBursts homepage (link) contains links to the various re-sources. Pre-built packages are provided for Windows, OS X and Linux. Installation instructions can be found in the Reference Documentation (link). A description of FRETBursts execution using Jupyter notebooks is reported in SI S1. Detailed information on development style, testing strategies and contributions guidelines are reported in SI S2. Finally, to facilitate evaluation and comparison with other software, we set up an on-line services allowing to execute FRETBursts without requiring any installation on the user’s computer (link).

**Table 1:** Photon selection syntax (non-ALEX)

| Photon selection | code |
| --- | --- |
| All-photons | `Ph_sel('all')` |
| D-emission | `Ph_sel(Dex='Dem')` |
| A-emission | `Ph_sel(Dex='Aem')` |

## 3 Architecture and Concepts

In this section, we introduce some general burst analysis concepts and notations used in FRETBursts.

### 3.1 Photon Streams

The raw data collected during a smFRET experiment consists in one or more arrays of photon timestamps, whose temporal resolution is set by the acquisition hardware, typically between 10 and 50 ns. In single-spot measurements, all timestamps are stored in a single array. In multispot measurements [42], there are as many timestamps arrays as excitation spots. Each array contains timestamps from both donor (D) and acceptor (A) channels. When alternating excitation lasers are used (ALEX measurements) [16], a further distinction between photons emitted during the D or A excitation periods can be made.

In FRETBursts, the corresponding sets of photons are called “photon streams” and are specified with a Ph_sel object (link). In non-ALEX smFRET data, there are 3 photon streams (table 1), while in two-color ALEX data, there are 5 streams (table 2).

The `Ph_sel` class (link) allows the specification of any combination of photon streams. For example, in ALEX measurements, the D-emission during A-excitation stream is usually ignored because it does not contain any useful signal [16]. To indicate all but photons in this photon stream, the syntax is `Ph_sel(Dex=’DAem’, Aex=’Aem’`), which indicates selection of donor and acceptor photons (`DAem`) during donor excitation (Dex) and only acceptor photons (`Aem`) during acceptor excitation (`Aex`).

### 3.2 Background Definitions

An estimation of the background rates is needed to both select a proper threshold for burst search, and to correct the raw burst counts by subtracting background counts.

**Table 2:** Photon selection syntax (ALEX)

| Photon selection | code |
| --- | --- |
| All-photons | `Ph_sel('all')` |
| D-emission during D-excitation | `Ph_sel(Dex='Dem')` |
| A-emission during D-excitation | `Ph_sel(Dex='Aem')` |
| D-emission during A-excitation | `Ph_sel(Aex='Dem')` |
| A-emission during A-excitation | `Ph_sel(Aex='Aem')` |

The recorded stream of timestamps is the result of two processes: one characterized by a high count rate, due to fluorescence photons of single molecules crossing the excitation volume, and another characterized by a lower count rate, due to “background counts” originating from detector dark counts, afterpulsing, out-of-focus molecules and sample scattering and/or impurities [20,48]. The signature of these two types of processes can be observed in the interphoton delays distribution (i.e. the waiting times between two subsequent timestamps) as illustrated in figure 1(a). The “tail” of the distribution (a straight line in semi-log scale) corresponds to exponentially-distributed time-delays, indicating that those counts are generated by a Poisson process. At short timescales, the distribution departs from an exponential due to the contribution of the higher rate process of single molecules traversing the excitation volume. To estimate the background rate (i.e. the inverse of the exponential time constant), it is necessary to define a time-delay threshold above which the distribution can be considered exponential. Finally, a parameter estimation method needs to be specified, such as Maximum Likelihood Estimation (MLE) or nonlinear least squares curve fitting of the time-delay histogram (both supported in FRETBursts).

It is advisable to monitor the background as a function of time throughout the measurement, in order to account for possible variations. Experimentally, we found that when the background is not constant, it usually varies on time scales of tens of seconds (see figure 2). FRETBursts divides the acquisition in constant-duration time windows called *background periods* and computes the background rates for each of these windows (see section 4.3). Note that FRETBursts uses these local background rates also during burst search, in order to compute time-dependent burst detection thresholds and for background correction of burst data (see section 4.4).

### 3.3 The Data Class

The Data class (link) is the fundamental data container in FRETBursts. It contains the measurement data and parameters (attributes) as well as several methods for data analysis (background estimation, burst search, etc…). All analysis results (bursts data, estimated parameters) are also stored as Data attributes.

There are 3 important “burst counts” attributes which contain the number of photons detected in the donor or the acceptor channel during donor or acceptor excitation periods (table 3). The attributes in table 3 are background-corrected by default. Furthermore, na is corrected for leakage and direct excitation (section 4.5) if the relative coefficients are specified (by default they are set to 0). There is also a closely related attribute named nda for donor photons detected during acceptor excitation. nda is normally neglected as it only contains background.

**Table 3:** Data attributes names and descriptions for burst photon counts in different photon streams.

| Name | Description |
| --- | --- |
| `nd` | number of photons detected by the donor channel (during donor excitation period in ALEX case) |
| `na` | number of photons detected by the acceptor channel (during donor excitation period in ALEX case) |
| `naa` | number of photons detected by the acceptor channel during acceptor excitation period (present only in ALEX measurements) |

**Python details** Many Data attributes are lists of arrays (or scalars) with the length of the lists equal to the number of excitation spots. This means that in single-spot measurements, an array of burst-data is accessed by specifying the index as 0, for example `Data.nd[0]`. Data implements a shortcut syntax to access the first element of a list with an underscore, so that an equivalently syntax is `Data.nd_` instead of `Data.nd[0]`.

### 3.4 Introduction to Burst Search

Identifying single-molecule fluorescence bursts in the stream of photons is one of the most crucial steps in the analysis of freely-diffusing single-molecule FRET data. The widely used “sliding window” algorithm, introduced by the Seidel group in 1998 [10, 12], involves searching for *m* consecutive photons detected during a period shorter than Δ*t*. In other words, bursts are regions of the photon stream where the local rate (computed using *m* photons) is above a minimum threshold rate. Since a universal criterion to choose the rate threshold and the number of photons *m* is, as of today, lacking, it has become a common practice to manually adjust those parameters for each specific measurement. Commonly employed values for m are between 5 and 15 photons.

A more general approach consists in taking into account the background rate of the specific measurements and in choosing a rate threshold that is *F* times larger than the background rate (typical values for *F* are between 4 and 9). This approach ensures that all resulting bursts have a signal-to-background ratio (SBR) larger than (*F* — 1) [46]. A consistent criterion for choosing the threshold is particularly important when comparing different measurements with different background rates, when the background significantly varies during measurements or in multi-spot measurements where each spot has a different background rate.

**Figure 1:**
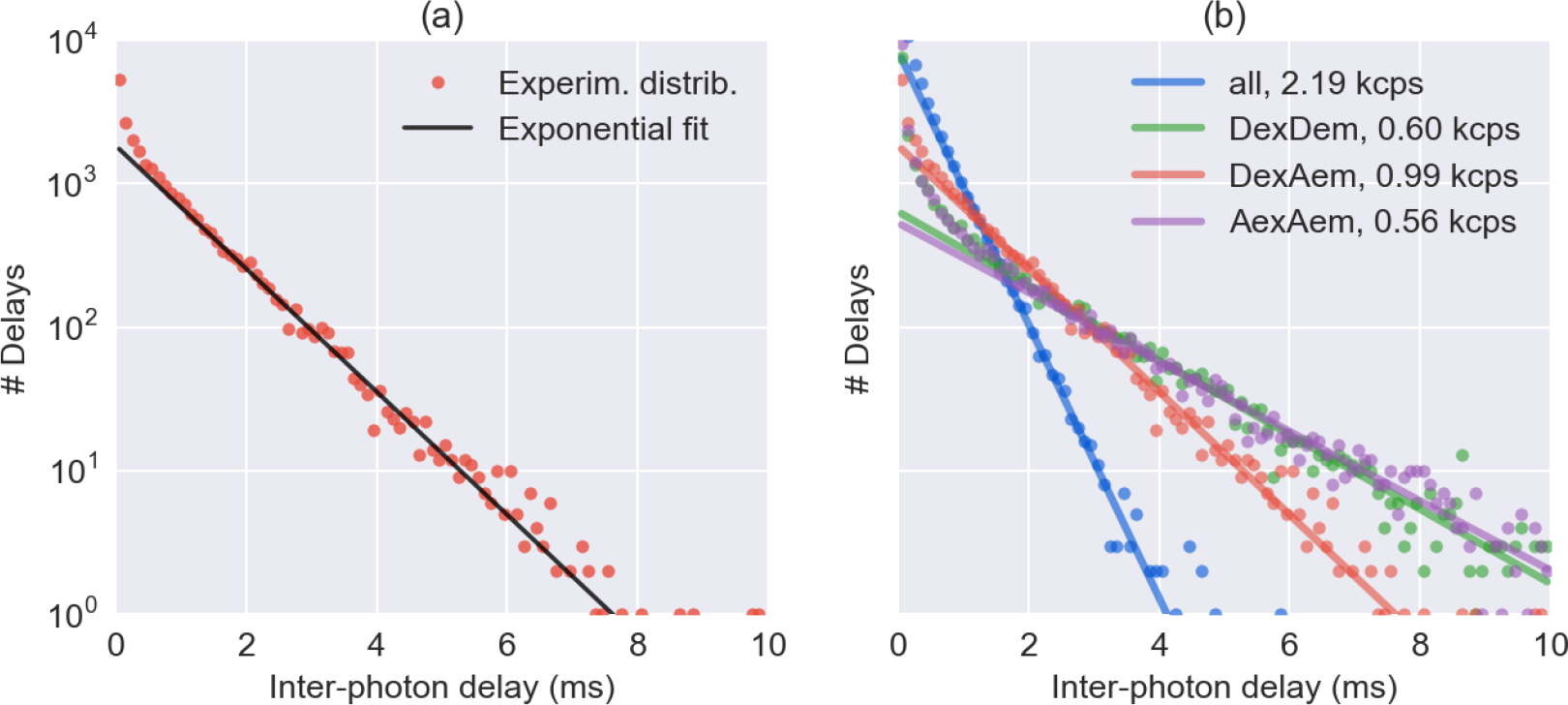
**Inter-photon delays fitted with and exponential function**. Experimental distributions of inter-photon delays (*dots*) and corresponding fits of the exponential tail (*solid lines*). (*Panel a*) An example of inter-photon delays distribution (*red dots*) and an exponential fit of the tail of the distribution (*black line*). (*Panel b*) Inter-photon delays distribution and exponential fit for different photon streams as obtained with `dplot`(`d, hist_bg`). The *dots* represent the experimental histogram for the different photon streams. The *solid lines* represent the corresponding exponential fit of the tail of the distributions. The legend shows abbreviations of the photon streams and the fitted background rates.

**Figure 2:**
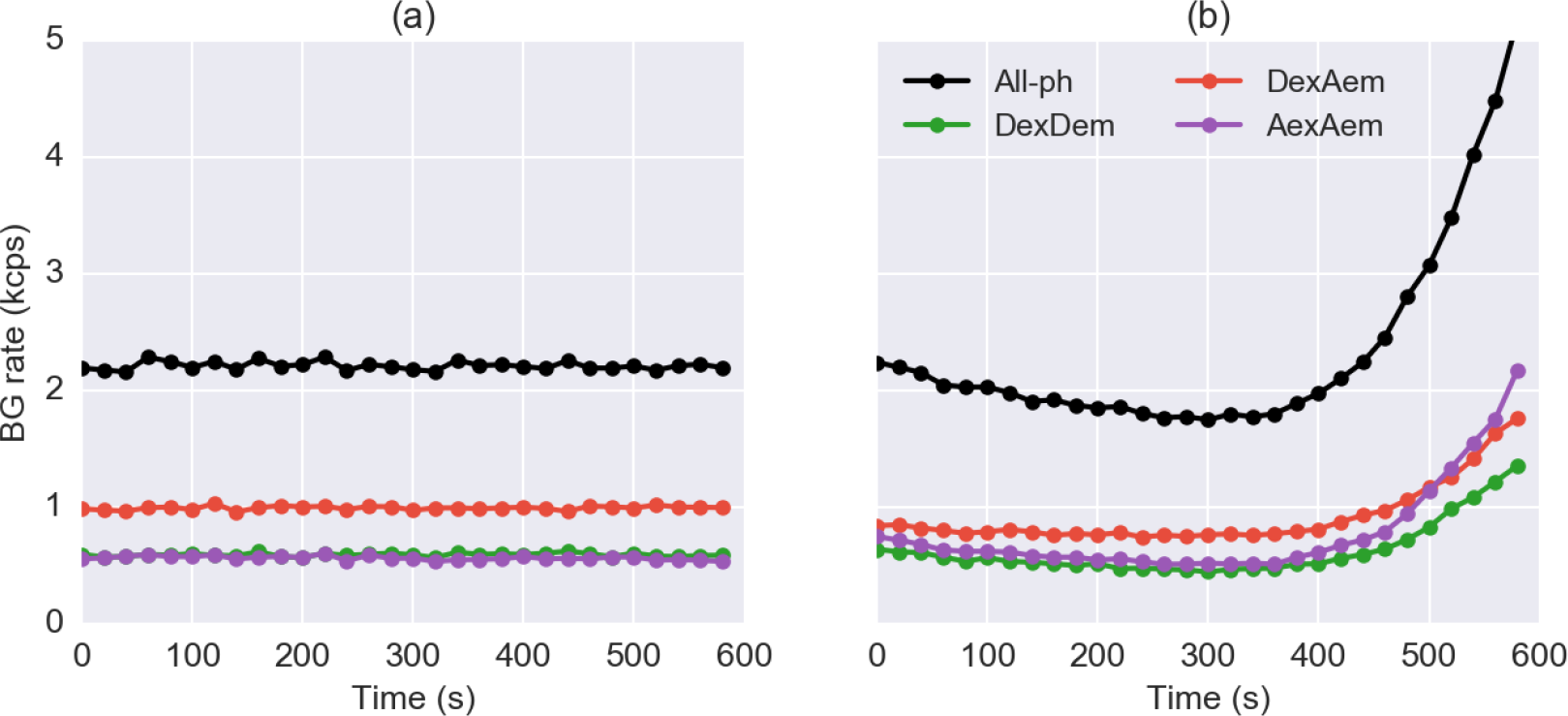
**Background rates as a function of time**. Estimated background rate as a function of time for two μs-ALEX measurements. Different colors represent different photon streams. (*Panel a*) A measurement performed with a sealed sample chamber exhibiting constant a background as a function of time. (*Panel b*) A measurement performed on an unsealed sample exhibiting significant background variations due to sample evaporation and/or photobleaching (likely impurities on the cover-glass). These plots are produced by the command `dplot`(`d, timetrace_bg`) after estimation of background. Each data point in these figures is computed for a 30 s time window.

A second important aspect of burst search is the choice of photon stream used to perform the search. In most cases, for instance when identifying FRET sub-populations, the burst search should use all photons, the so called all-photon burst search (APBS) [10,12,17]. In other cases, for example when focusing on donor-only or acceptor-only populations, it is better to perform the search using only donor or acceptor signal. In order to handle the general case and to provide flexibility, FRETBursts allows performing the burst search on arbitrary selections of photons (see section 3.1 for more information on photon stream definitions).

Additionally, Nir *et al.* [17] proposed a dual-channel burst search (DCBS) which can help mitigating artifacts due to photophysics effects such as blinking. During DCBS, a search is performed on two photon streams and bursts are defined as periods during which both photon streams exhibit a rate higher than the threshold, implementing the equivalent of an AND logic operation. Conventionally, the term DCBS refers to a burst search where the two photon streams are (1) all photons during donor excitation (`Ph_sel`(`Dex=’DAem’`)) and (2) acceptor channel photons during acceptor excitation (`Ph_sel`(Aex=’Aem’)). In FRETBursts, the user can choose arbitrary photon streams as input, an in general this kind of search is called a “AND-gate burst search”. For additional details on burst search refer to the documentation(link).

After burst search, it is necessary to further select bursts, for instance by specifying a minimum number of photons (or burst size). In the most basic form, this selection can be performed during burst search by discarding bursts with size smaller than a threshold *L* (typically 30 or higher), as originally proposed by Eggeling *et al.* [10]. This method, however, neglects the effect of background and γ factor on the burst size and can lead to a selection bias for some channels and/or sub-populations. For this reason, we suggest performing a burst size selection after background correction, taking into account the γ factor, as discussed in sections 3.5 and 4.6. In special cases, users may choose to replace (or combine) the burst selection based on burst size with another criterion such as burst duration or brightness (see section 4.6).

### 3.5 Corrected Burst Sizes and Weights

The number of photons detected during a burst-the “burst size” -is computed using either all photons, or photons detected during donor excitation period. To compute the burst size, FRETBursts uses one of the following formulas:

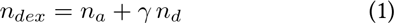

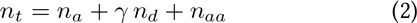

where *n_d_*, *n_a_* and *n_aa_* are, similarly to the attributes in table 3, the background-corrected burst counts in different channels and excitation periods. The factor γ takes into account different fluorescence quantum yields of donor and acceptor fluorophores and different photon detection efficiencies between donor and acceptor detection channels [16,49]. Eq. 1 includes counts collected during donor excitation periods only, while eq. 2 includes all counts. Burst sizes computed according to eq. 1 or 2 are called γ-corrected burst sizes.

The burst search algorithm yields a set of bursts whose sizes approximately follow an exponential distribution. Compared to bursts with smaller sizes, bursts with large sizes are less frequent, but contain more information per-burst (having higher SNR). Therefore, selecting bursts by size is an important step (see section 4.6). A threshold set too low may result in unresolvable sub-populations because of broadening of FRET peaks and appearance of shot-noise artifacts in the FRET (and *S*) distribution (i.e. spurious narrow peaks due to *E* and *S* being computed as the ratio of small integers). Conversely, too large a threshold may result in too low a number of bursts therefore poor representation of the FRET distribution. Additionally, especially when computing fractions of sub-populations (e.g. ratio of number of bursts in each sub-population), it is important to use γ-corrected burst sizes as selection criterion, in order to avoid underrepresenting some FRET sub-populations due to different quantum yields of donor and acceptor dyes and/or different photon detection efficiencies of donor and acceptor channels.

An alternative method to apply γ correction consist in discarding a constant fraction of photons chosen randomly from either the D_em_ or A_em_ photon stream [17]. This simple method transforms the measurement data in order to achieve γ = 1, overcoming the issue of selection bias between populations. This approach has also the advantage of preserving the binomial distribution of D and A photons in each burst, so that peaks of FRET populations are easier to model statistically. The only drawback is that, by discarding a fraction of photons, this method leads to information loss and therefore to a potential decrease in sensitivity and/or accuracy.

A simple way to mitigate the dependence of the FRET distribution on the burst size selection threshold is weighting bursts proportionally to their size so that the bursts with largest sizes will have the largest weights. Using size as weights (instead of any other monotonically increasing function of size) can be justified noticing that the variance of bursts proximity ratio (PR) is inversely proportional to the burst size (see SI S6 for details).

In general, a weighting scheme is used for building efficient estimators for a population parameter (e.g. the population FRET efficiency *E_p_*). But, it can also be used to build weighted histograms or Kernel Density Estimation (KDE) plots which emphasize FRET subpopulations peaks without excluding small size bursts. Traditionally, for optimal results when not using weights, the FRET histogram is manually adjusted by finding an ad-hoc (high) size-threshold which selects only bursts with the highest size (and thus lowest variance). Building size-weighted FRET histograms is a simple method to balance the need of reducing the peaks width with the need of including as much bursts as possible to reduce statistical noise. As a practical example, by fixing the burst size threshold to a low value (e.g. 10-20 photons) and using weights, is possible to build a FRET histogram with well-defined FRET sub-populations peaks without the need of searching an optimal burst-size threshold (SI S6).

**Python details** FRETBursts has the option to weight bursts using γ-corrected burst sizes which optionally include acceptor excitation photons naa. A weight proportional to the burst size is applied by passing the argument `weights`=’`size`’ to histogram or KDE plot functions. The weights keyword can be also passed to fitting functions in order to fit the weighted E or S distributions (see section 4.7). Other weighting functions (for example depending quadratically on the size) are listed in the `fret_fit.get_weights` documentation (link). However, using weights different from the size is not recommended due to their less efficient use of burst information (SI S6).

## 4 smFRET Burst Analysis

### 4.1 Loading Data

While FRETBursts can load several data files formats, we encourage users to adopt the recently introduced Photon-HDF5 file format [40]. photon-HDF5 is anHDF5-based, open format, specifically designed for freely-diffusing smFRET and other timestamp-based experiments. Photon-HDF5 is a self-documented, platform-and language-independent binary format, which supports compression and allows saving photon data (e.g. timestamps) and measurement-specific metadata (e.g. setup and sample information, authors, provenance, etc.). Moreover, Photon-HDF5 is designed for longterm data preservation and aims to facilitate data sharing between different software and research groups. All example data files provided with FRETBursts use the Photon-HDF5 format.

To load data from a Photon-HDF5 file, we use the function loader.photon_hdf5 (link):

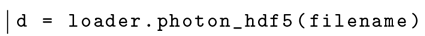

where filename is a string containing the file path. This command loads the measurement data into the variable d, a Data object (see section 3.3).

The same command can load data from a variety of smFRET measurements supported by the Photon-HDF5 format, taking advantage of the rich metadata included with each file. For instance, data generated using different excitation schemes such as CW excitation or pulsed excitation, single-laser vs two alternating lasers, etc., or with any number of excitation spots, are automatically recognized and interpreted accordingly.

FRETBursts also supports loading μs-ALEX data stored in .sm files (a custom binary format used in the Weiss lab) and ns-ALEX data stored in .spc files (a binary format used by TCSPC Becker & Hickl acquisition hardware). Alternatively, these and other formats (such as ht3, a binary format used by PicoQuant hardware) can be converted into Photon-HDF5 files using phconvert, a file conversion library and utility for Photon-HDF5 (link). More information on loading different file formats can be found in the loader module’s documentation (link).

### 4.2 Alternation Parameters

For μs-ALEX and ns-ALEX data, Photon-HDF5 normally stores parameters defining alternation periods corresponding to donor and acceptor laser excitation. At load time, a user can plot these parameters and change them if deemed necessary. In μs-ALEX measurements [50], CW laser lines are alternated on timescales of the order of 10 to 100 μs. Plotting an histogram of timestamps modulo the alternation period, it is possible to identify the donor and acceptor excitation periods (see figure 3a). In ns-ALEX measurements [44], pulsed lasers with equal repetition rates are delayed with respect to one another with typical delays of 10 to 100 ns. In this case, forming an histogram of TCSPC times (nanotimes) will allow the definition of periods of fluorescence after excitation of either the donor or the acceptor (see figure 3b). In both cases, the function plot_alternation_hist (link) will plot the relevant alternation histogram (figure 3) using currently selected (or default) values for donor and acceptor excitation periods.

To change the period definitions, we can type:

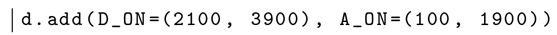

where `D_ON` and `A_ON` are pairs of numbers (tuples in Python) representing the *start* and *stop* values for D or A excitation periods. The previous command works for both μs-ALEX and ns-ALEX measurements. After changing the parameters, a new alternation plot will show the updated period definitions.

The alternation period definition can be applied to the data using the function loader.`alex_apply_period(d)` (link):

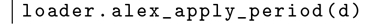

After this command, d will contain only photons inside the defined excitation periods. If the user needs to update the periods definition, the data file will need to be reloaded and the steps above repeated as described.

### 4.3 Background Estimation

The first step of smFRET analysis involves estimating background rates. For example, the following command:

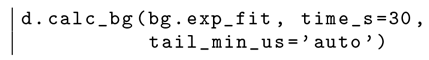

estimates the background rates in windows of30 s using the default iterative algorithm for choosing the fitting threshold (section 3.2). Beginner users can simply use the previous command and proceed to burst search (section 4.4). For more advanced users, this section provides details on the different background estimation and plotting functions provided by FRETBursts.

**Figure 3:**
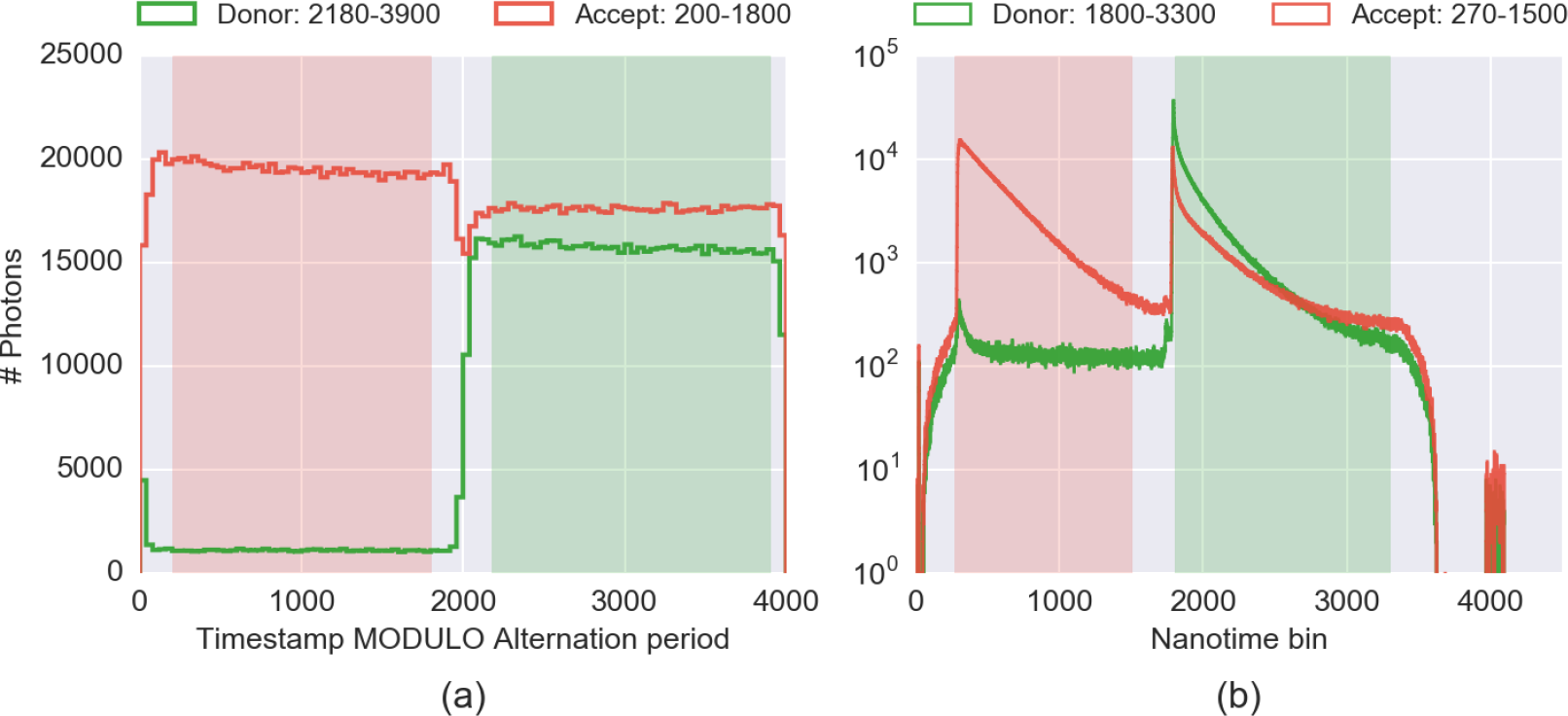
**Alternation histograms for μs-ALEX and ns-ALEX measurements**. Histograms used for the selection/determination of the alternation periods for two typical smFRET-ALEX experiments. Distributions of photons detected by donor channel are in green, and by acceptor channel in *red*. light *green* and *red* shaded areas indicate the donor and acceptor period de.nitions. (a) μs-ALEX alternation histogram, i.e. histogram of timestamps modulo the alternation period for a smFRET measurement (in timestamp clock unit). (b) ns-ALEX TCSPC nanotime histogram for a smFRET measurement (in TDC or TAC bin unit). Both plots have been generated by the same plot function (`plot alternation hist`()).
Additional information on these speci.c measurements can be found in the a.ached notebook (link).

First, we show how to estimate the background every 30 s, using a fixed inter-photon delay threshold of 2 ms (the same for all photon streams):

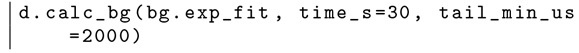

The first argument (`bg.exp_fit`) is the function used to fit the background rate for each photon stream (see section 3.2). The function `bg.exp_fit` estimates the background using a maximum likelihood estimation (MLE) of the delays distribution. The second argument, `time_s`, is the duration of the *background period* (section 3.2) and the third, `tail_min_us`, is the minimum inter-photon delay to use when fitting the distribution to the specified model function. To use different thresholds for each photon stream we pass a tuple (i.e. a comma-separated list of values, link) instead of a scalar. However, the recommended approach is to choose the threshold automatically using `tail_min_us`=’`auto`’. This approach uses an heuristic algorithm described in the *Background estimation* section of the μs-ALEX tutorial (link). Finally, it is possible to use a rigorous but slower approach to find an optimal threshold, as described in SI S5.

FRETBursts provides two kinds of plots to represent the background. One shows the histograms of inter-photon delays compared to the fitted exponential distribution, shown in figure 1) (see section 3.2 for details on the inter-photon distribution). This plot is created with the command:

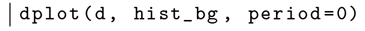

This command illustrates the general form of a plotting commands in FRETBursts, as described in SI S4. Here we only note that the argument period is an integer specifying the background period to be plotted (when omitted, the default is 0, i.e. the first period). Figure 1 allows to quickly identify pathological cases where the background fitting procedure returns unreasonable values.

The second background-related plot represents a timetrace of background rates, as shown in figure 2. This plot allows monitoring background rate variations occurring during the measurement and is obtained with the command:

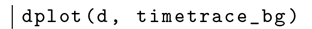

Normally, samples should have a fairly constant background rate as a function of time as in figure 2(a). However, sometimes, non-ideal experimental conditions can yield a time-varying background rate, as illustrated in figure 2(b). A possible reason for the observed behavior could be buffer evaporation from an open sample (we strongly recommend using a sealed observation chamber whenever possible). Additionally, cover-glass impurities can contribute to the background. These impurities tend to bleach on timescales of minutes resulting in background variations during the course of the measurement.

**Python details** The estimated background rates are stored in the Data attributes `bg_dd`, `bg_ad` and `bg_aa`, corresponding to photon streams `Ph_sel`(`Dex`=’`Dem`’), `Ph_sel`(`Dex`=’`Aem’`) and `Ph_sel`(`Aex`=’`Aem`) respectively. These attributes are lists of arrays (one array per excitation spot). The arrays contain the estimated background rates in the different time windows (background periods). Additional background fitting functions (e.g. least-square fitting of interphoton delay histogram) are available in bg namespace (i.e. the background module, link).

### 4.4 Burst Search

Following background estimation, burst search is the next step of the analysis. In FRETBursts, a standard burst search using a single photon stream (see section 3.4) is performed by calling the Data.burst_search method (link). For example, the following command:

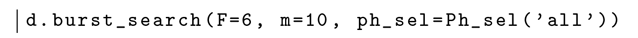

performs a burst search on all photons (`ph_sel`’`Ph_sel`(’`all`’)), with a count rate threshold equal to 6 times the local background rate (`f=6`), using 10 consecutive photons to compute the local count rate (`m=10`). A different photon stream, threshold (*F*) or number of photons m can be selected by passing different values. These parameters are good general-purpose starting point for smFRET analysis but they can be adjusted if needed.

Note that the previous burst search does not perform any burst size selection (however, by definition, the minimum bursts size is effectively *m*). An additional parameter *L* can be passed to impose a minimum burst size before any correction. However, it is recommended to select bursts only after applying background corrections, as discussed in the next section 4.6.

It might sometimes be useful to specify a fixed photon-rate threshold, instead of a threshold depending on the background rate, as in the previous example. In this case, instead of *F*, the argument `min_rate_cps` can be used to specify the threshold (in counts-per-second). For example, a burst search with a 50 kcps threshold is performed as follows:

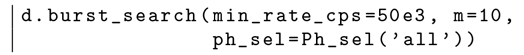

Finally, to perform a DCBS burst search (or in general an AND gate burst search, see section 3.4) we use the function `burst_search_and_gate` (link), as illustrated in the following example:

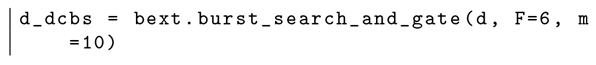

The last command puts the burst search results in a new copy of the `Data` variable `d` (in this example the copy is called `d_dcbs`). Since FRETBursts shares the timestamps and detectors arrays between different copies of Data objects, the memory usage is minimized, even when several copies are created.

**Python details** Note that, while `d.burst_search()` is a method of `Data, bext.burst_search_and_gate()` is a function in the `bext` module taking a `Data` object as a first argument and returning a new `Data object`.

The function burst_search_and_gate accepts optional arguments, `ph_sel1` and `ph_sel2`, whose default values correspond to the classical DCBS photon stream selection (see section 3.4). These arguments can be specified to select different photon streams than those used in a classical DCBS.

The bext module (link) collects “plugin” functions that provides additional algorithms for processing Data objects.

### 4.5 Bursts Corrections

In μs-ALEX, there are 3 important correction parameters: γ-factor, donor leakage into the acceptor channel and acceptor direct excitation by the donor excitation laser [16]. These corrections can be applied to burst data by simply assigning values to the respective Data attributes:

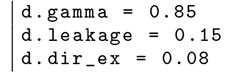

These attributes can be assigned either before or after the burst search. In the latter case, existing burst data are automatically updated using the new correction parameters.

These correction factors can be used to display corrected FRET distributions. However, when the goal is to fit the FRET efficiency of sub-populations, it is simpler to fit the background-corrected PR histogram and then correct the population-level PR value (see SI in [16]). Correcting PR of each population (instead of correcting the data in each burst) avoids distortion of the FRET distribution and keeps peaks of static FRET subpopulations closer to the ideal binomial statistics [19].

FRETBursts implements the correction formulas for *E* and *S* in the functions `fretmath.correct_E_gamma_leak_dir` and `fretmath.correct_S` (link). A derivation of these correction formulas (using computer-assisted algebra) can be found online as an interactive notebook (link).

### 4.6 Burst Selection

After burst search, it is common to select bursts according to different criteria. One of the most common is burst size.

For instance, to select bursts with more than 30 photons detected during the donor excitation (computed after background correction), we use following command:

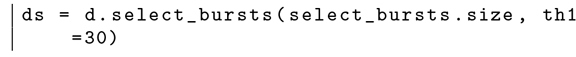

The previous command creates a new Data variable (ds) containing the selected bursts. th1 defines the lower bound for burst size, while `th2` defines the upper bound (when not specified, as in the previous example, the upper bound is +∞). As before, the new object (`ds`) will share the photon data arrays with the original object (`d`) in order to minimize the amount of used memory.

The first argument of `select_bursts` (link) is a python function implementing the “selection rule” (`select_bursts.size` in this example); all remaining arguments (only `th1` in this case) are parameters of the selection rule. The `select_bursts module` (link) contains numerous built-in selection functions (link). For example, `select_bursts.ES` is used to select a region on the E-S ALEX histogram, `select_bursts.width` to select bursts based on their duration. New custom criteria can be readily implemented by defining a new selection function, which requires only a couple of lines of code (see the `select_bursts` module’s source code for examples, link).

Finally, different criteria can be combined sequentially. For example, with the following commands:

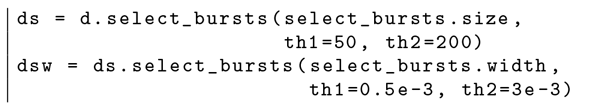

bursts in dsw will have sizes between 50 and 200 photons, and duration between 0.5 and 3 ms.

**Burst Size Selection** In the previous section, we selected bursts by size, using only photons detected in both D and A channels during D excitation (i.e. Dex photons), as in eq. 1. Alternatively, a threshold on the burst size computed including all photons can be applied by adding *n_aa_* to the burst size (see eq. 2). This is achieved by passing add_naa=True to the selection function. The complete selection command is:

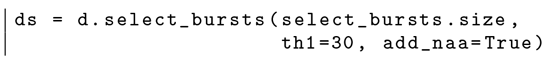

The result of this selection is plotted in figure 4. When add_naa is not specified, as in the previous section, the default is add_naa=False (i.e. compute size using only Dex photons).

Another important parameter for defining the burst size is the γ-factor. As noted in section 3.5, the γ-factor is used to compensate for different fluorescence quantum yields for the D and A fluorophores as well as different photon-detection efflciencies for the D and A channels. When y is significantly different from 1, neglecting its effect on burst size leads to over-representing one FRET population versus the others.

When the γ factor is known (and ≠ 1), a more unbiased selection of different FRET populations can be achieved passing the argument gamma to the selection function:

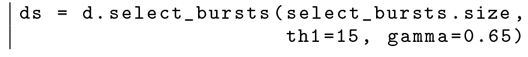

When not specified, γ = 1 is assumed. For more details on burst size selection, see the select_bursts.size documentation (link).

**Python details** The method to compute γ-corrected burst sizes (with or without addition of naa) is Data.burst_sizes (link).

**Select the FRET Populations** In smFRET-ALEX experiments, donor-only (D-only) and acceptor-only (A-only) populations can be detected in addition to the FRET population(s). In most cases, the D-only and A-only populations are of no interest and need to be filtered out.

In principle, using the E-S representation, D-only and A-only bursts can be excluded by selecting bursts within a range of *S* values (e.g. S=0.2-0.8). This approach, however, simply truncates the burst distribution with arbitrary thresholds and is therefore not recommended for quantitative assessment of FRET populations.

**Figure 4:**
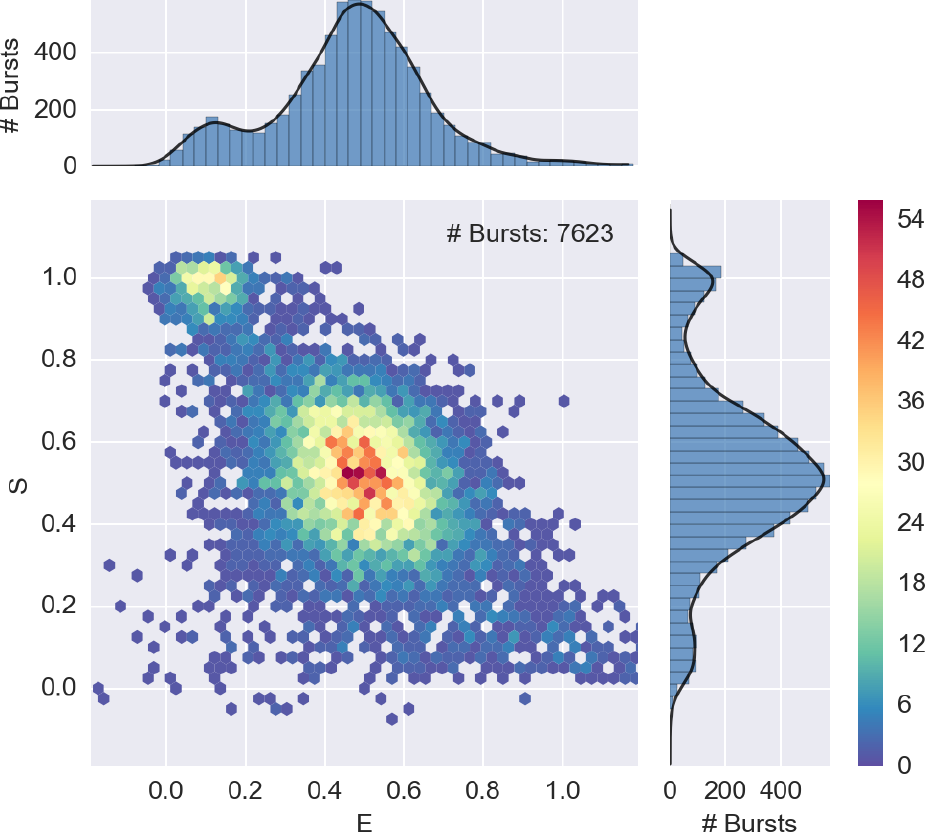
**E-S histogram showing FRET, D-only and A-only populations**. A 2-D ALEX histogram and marginal E and S histograms for a 40-bp dsDNA with D-A distance of 17 bases (Donor dye: ATT0550, Acceptor dye: ATT0647N). Bursts are selected with a size-threshold of 30 photons, including Aex photons. The plot is obtained with `alex_jointplot(ds)`. The 2D E-S distribution plot (join plot) is an histogram with hexagonal bins, which reduce the binning artifacts (compared to square bins) and naturally resembles a scatter-plot when the burst density is low (see SI S4). Three populations are visible: FRET population (middle), D-only population (top left) and A-only population (bottom, *S* < 0.2). Compare with figure 5 where the FRET population has been isolated.

**Figure 5:**
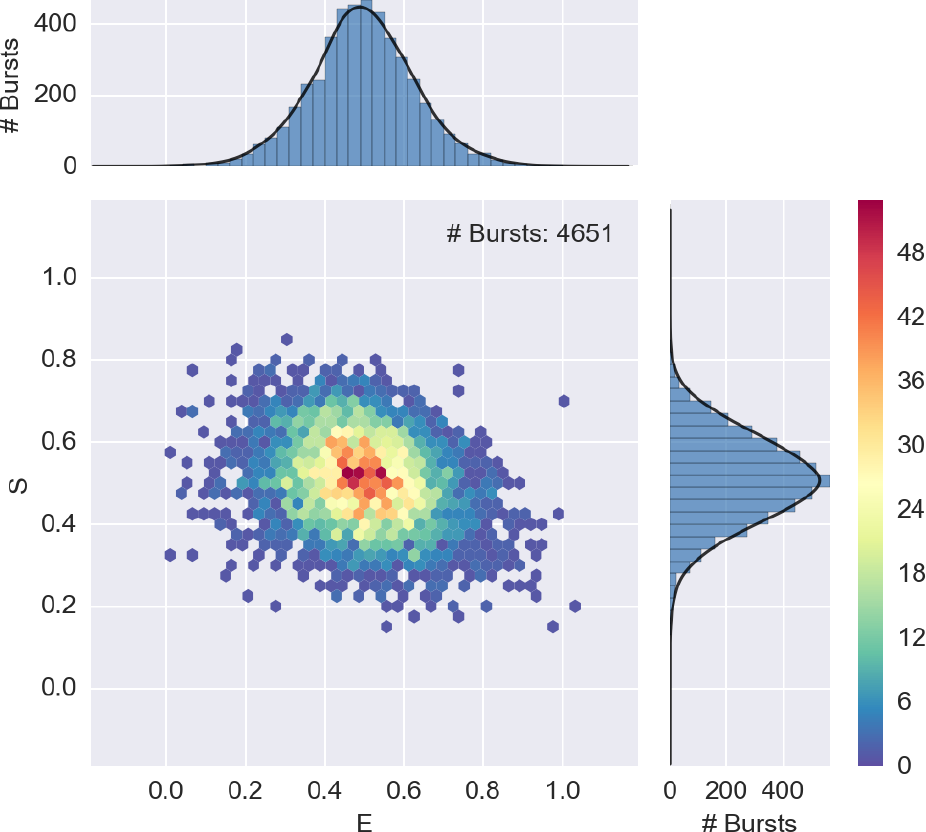
**E-S histogram after filtering out D-only and A-only populations**. 2-D ALEX histogram after selection of FRET population using the composition of two burst selection filters: (1) selection of bursts with counts in Dex stream larger than 15; (2) selection of bursts with counts in AexAem stream larger than 15. Compare to figure 4 where all burst populations (FRET, D-only and A-only) are reported.

An alternative approach consists in applying two selection filters sequentially. First, the A-only population is filtered out by applying a threshold on the number of photons during D excitation (Dex). Second, the D-only population is filtered out by applying a threshold on the number of A photons during A excitation (AexAem). The commands for these combined selections are:

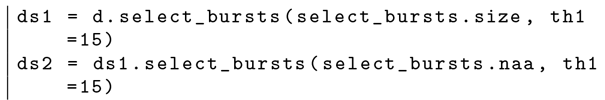

Here, the variable ds2 contains the combined burst selection. Figure 5 shows the resulting filtered FRET population obtained with the previous selection.

### 4.7 Population Analysis

Typically, after bursts selection, E or S histograms are fitted to a model. FRETBursts `mfit` module allows fitting histograms of bursts quantities (i.e. E or S) with arbitrary models. In this context, a model is an object specifying a function, the parameters varied during the fit and optional constraints for these parameters. This concept of model is taken from *lmfit* [51], the underlying library used by FRETBursts to perform the fits.

Models can be created from arbitrary functions. FRETBursts includes predefined (i.e. built-in) models such as 1- to 3-Gaussian peaks or 2-Gaussian connected by a flat “plateau”. The latter is an empirical model which can be used to more accurately fit the center values of two populations when the peaks are connected by intermediate-FRET bursts (for the analytical definition of this function see the documentation, link). Built-in models are created by calling a corresponding factory function (whose names start with `mfit.factory_`) which initializes the parameters with values and constraints suitable for E and S histograms fits (see *Factory Functions* documentation, link).

As an example, we can fit the E histogram of bursts in the ds variable with two Gaussian peaks with the following command:

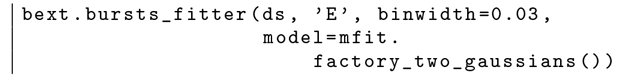

Changing ‘`E`’ with ‘`S`’ will fit the S histogram instead. The binwidth argument specifies the histogram bin width and the model argument defines which model shall be used for fitting.

All fitting results (including best fit values, uncertainties, etc.), are stored in the E_fitter (or S_fitter) attributes of the Data variable (named ds here). To print a comprehensive summary of the fit results, including uncertainties, reduced *χ^2^* and correlation between parameters, we can use the following command:

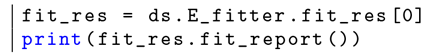

Finally, to plot the fitted model together with the FRET histogram, as shown in figure 6, we pass the parameter show_model=True to the hist_fret function (see section S4 for an introduction to plotting in FRETBursts):

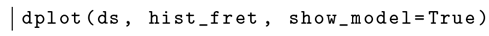

For more examples on fitting bursts data and plotting results, refer to the fitting section of the μs-ALEX notebook (link), the *Fitting Framework* section of the documentation (link) as well as the documentation for `bursts_fitter` function (link).

**Python details** Models returned by FRETBursts’s factory functions (`mfit.factory_*`) are lmfit.Model objects (link). Custom models can be created by calling `lmfit.Model` directly. When an `lmfit.Model` is fitted, it returns a `ModelResults` object (link), which contains all information related to the fit (model, data, parameters with best values and uncertainties) and useful methods to operate on fit results. FRETBursts puts a ModelResults object of each excitation spot in the list `ds.E_fitter.fit_res`. For instance, to obtain the reduced χ^2^ value of the E histogram fit in a single-spot measurement d, we use the following command:

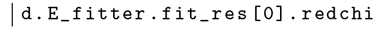

**Figure 6:**
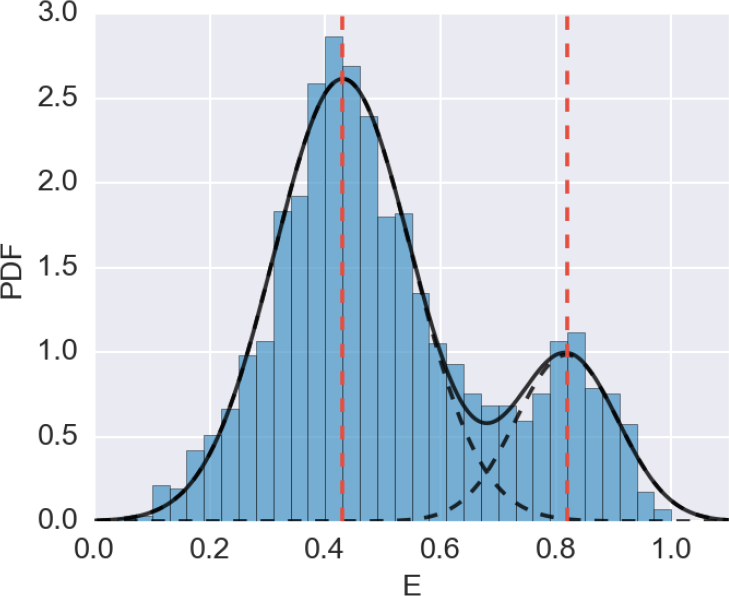
**FRET histogram fitted with two Gaussians**.Example of a FRET histogram fitted with a 2-Gaussian model. After performing the fit (see main text), the plot is generated with `dplot(ds, hist_fret, show_model=True`).

Other useful attributes are aic and bic which contain statistics for the Akaike information criterion (AIC) [52] and the Bayes Information criterion (BIC) [53]. AIC and BIC are general-purpose statistical criteria for comparing the suitability of multiple non-nested models according to the data. By penalizing models with higher number of parameters, these criteria strike a balance between the need of achieving high goodness of fit with the need of keeping the model complexity low to avoid overfitting.

Examples of definition and modification of fit models are provided in the aforementioned μs-ALEX notebook (link). Users can also refer to the comprehensive lmfit’s documentation (link).

### 4.8 FRET Dynamics

Single-molecule FRET histograms show more information than just mean FRET efficiencies. While in general the presence of several peaks clearly indicates the existence of multiple subpopulations, a single peak cannot a priori be associated with a single population defined by a unique FRET efficiency without further analysis.

Shot-noise analysis [17] or probability distribution analysis (PDA) [18, 54] allow to compute the minimum width of a static FRET population (i.e. caused by the statistics of discrete photon-detection events). Typically, several mechanisms contribute to the broadening ofthe experimental FRET peak beyond the shot-noise limit. These include heterogeneities in the sample resulting in a distribution of Förster radii, or actual conformational changes giving rise to a distribution of D-A distances [8].

Gopich and Szabo developed an elegant analytical model for the FRET distribution of *M* interconverting states based on superposition of Gaussian peaks [55]. Unfortunately, the method is not of straightforward application for freely-diffusing data as it requires a special selection criterion for filtering bursts with quasi-Poisson rates. Santoso et al. [56] and Kalinin et al. [57] extended the PDA approach to estimate conversion rates between different states by comparing FRET histograms as a function of the time-bin size. In addition, GopichandSzabo [58,59] developedarelatedmethodto compute conversion rates using a likelihood function which depends on photon timestamps (overcoming the time binning and FRET histogramming step and directly applicable to freely-diffusing data). In case of measurement including lifetime, the multiparameter fluorescence detection (MFD) method allows to identify dynamics from the deviation from the linear relation between lifetime and E [8]. Hoffman et al. [60] proposed a method called RASP (recurrence analysis of single particles) to extend the timescale of detectable kinetics. Hoffman et al. compute the probability that two nearby bursts are due to the same molecule and therefore allows setting a time-threshold for considering consecutive bursts as the same single-molecule event.

Other interesting approaches include combining smFRET and FCS for detecting and quantifying kinetics on timescales much shorter than the diffusion time [61–63]. In addition, Bayes-based methods have been proposed to fit static populations [64, 65], or to study dynamics [66].

Finally, two related methods for discriminating between static heterogeneity and sub-millisecond dynamics are Burst Variance Analysis (BVA) proposed by Torella et al. [23] and two-channel kernel density estimator (2CDE) proposed by Tomov et al. [24]. The BVA method is described in the next section. The 2CDE method, which has been implemented in FRETBursts, computes local photon rates from timestamps within bursts using Kernel Density Estimation (KDE) (FRETBursts includes general-purpose functions to compute KDE of photon timestamps in the `phrates` module, (link)). From time variations of local rates, it is possible to infer the presence of some dynamics. In particular, the 2CDE method builds, for each burst, a quantity (*E*)*D* (or (1 — *E*)*A*), which is equal to the burst average E when no dynamics is present, but is biased toward an higher (or lower) value in presence of dynamics. From these quantities, a burst “estimator” (called FRET-2CDE) is derived. Fora user, the 2CDE method amounts to plotting the 2-D histogram of *E* versus FRET-2CDE, and assessing the vertical position of the various populations: populations centered around FRET-2CDE=10 undergo no dynamics, while population biased towards higher FRET-2CDE values undergo dynamics.

The BVA and 2CDE methods are implemented in two notebooks included with FRETBursts (BVA link, 2CDE link). To use them, a user needs to download the relevant notebook and run the anaysis therein. The other methods mentioned in this section are not currently implemented in FRETBursts. However, users can easily implement any additional methods in FRETBursts, using its built-in burst analysis and timestamps/bursts manipulation functions. In the next section, we show how to perform low-level analysis of timestamps and bursts data by implementing the BVA method from scratch. An additional example showing how to split bursts in constant time-bins can be found in the respective FRETBursts notebook (link). These examples serve as a guide for implementing new methods. We welcome researchers willing to implement new methods to ask questions on GitHub or on the mailing list. We also encourage sharing eventual new methods implemented in FRETBursts for the benefit the entire community.

## 5 Implementing Burst Variance Analysis

In this section, we describe how to implement burst variance analysis (BVA) as described in [23]. FRETBursts provides well-tested, general-purpose functions for timestamps and burst data manipulation and therefore simplifies implementing custom burst analysis algorithms such as BVA.

### 5.1 BVA Overview

The BVA method has been developed to identify the presence of dynamics in FRET distributions [23], and has been successfully applied to identify biomolecular processes with dynamics on the millisecond time-scale [23,67].

The basic idea behind BVA is to subdivide bursts into contiguous burst chunks (sub-bursts) comprising a fixed number *n* of photons, and to compare the empirical variance of acceptor counts of all sub-bursts in a burst, with the theoretical shot-noise-limited variance. An empirical variance of sub-bursts larger than the shot-noise-limited value indicates the presence of dynamics.

In a FRET (sub-)population originating from a single static FRET efflciency, the sub-bursts acceptor counts *n_a_* can be modeled as a binomial-distributed random variable *N_a_* ∼ B(*n*, *E_p_*), where *n* is the number of photons in each sub-burst and *E_p_* is the estimated population proximity-ratio (PR). Note that we can use the PR because, regardless of the molecular FRET efflciency, the detected counts are partitioned between donor and acceptor channels according to a binomial distribution with success probability equal to the PR. The only approximation done here is neglecting the presence of background (a reasonable approximation since the backgrounds counts are in general a very small fraction of the total counts). We refer the interested reader to [23] for further discussion.

If *N_a_* follows a binomial distribution, the random variable *E*_sub_ = *N_a_/n*, has a standard deviation reported in eq. 3.

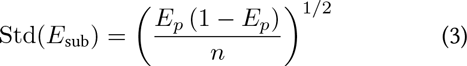

BVA analysis consists of four steps: 1) dividing bursts into consecutive sub-bursts containing a constant number of consecutive photons *n*, 2) computing the PR of each sub-burst, 3) calculating the empirical standard deviation (*s_E_*) of sub-bursts PR in each burst, and 4) comparing *s_E_* to the expected standard deviation of a shot-noise-limited distribution (eq. 3). If, as in figure 7, the observed FRET efflciency distribution originates from a static mixture of sub-populations (of different non-interconverting molecules) characterized by distinct FRET efflciencies, *s_E_* of each burst is only affected by shot-noise and will follow the expected standard deviation curve based on eq. 3. Conversely, if the observed distribution originates from biomolecules belonging to a single species, which interconverts between different FRET subpopulations (over times comparable to the diffusion time), as in figure 8, *s_E_* of each burst will be larger than the expected shot-noise-limited standard deviation, and will be located above the shot-noise standard deviation curve (right panel of figure 8).

### 5.2 BVA Implementation

The following paragraphs describe the low-level details involved in implementing the BVA using FRETBursts. The main goal is to illustrate a real-world example of accessing and manipulating timestamps and burst data. For a ready-to-use BVA implementation users can refer to the corresponding notebook included with FRETBursts (link).

**Python details** For BVA implementation, two photon streams are needed: all-photons during donor excitation (Dex) and acceptor photons during donor excitation (DexAem). These photon stream selections are obtained by computing boolean masks as follows (see section S3):

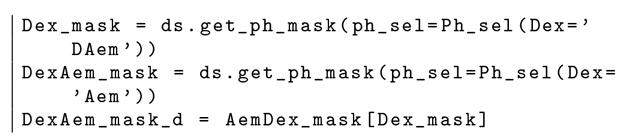

Here, the first two variables (`Dex_mask and DexAem_mask`) select photon from the all-photons timestamps array, while `DexAem_mask_d`, selects A-emitted photons from the array of photons emitted during D-excitation. As shown below, the latter is needed to count acceptor photons in burst chunks.

Next, we need to express bursts start-stop data as indexes of the D-excitation photon stream (by default burst start-stop indexes refer to all-photons timestamps array):

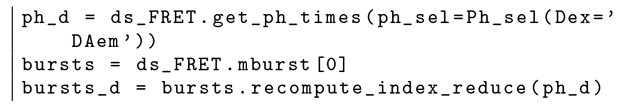

Here, `ph_d` contains the Dex timestamps, `bursts` the original burst data and `bursts_d` the burst data with start-stop indexes relative to `ph_d`.

Finally, with the previous variables at hand, the BVA algorithm can be easily implemented by computing the *s_E_* quantity for each burst:

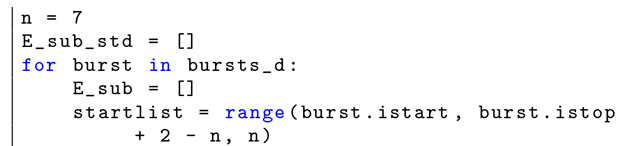

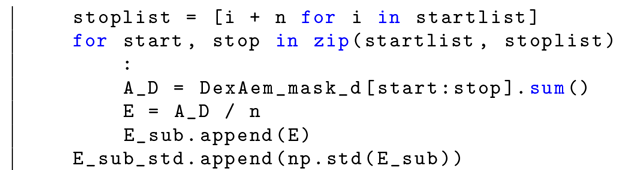

**Figure 7:**
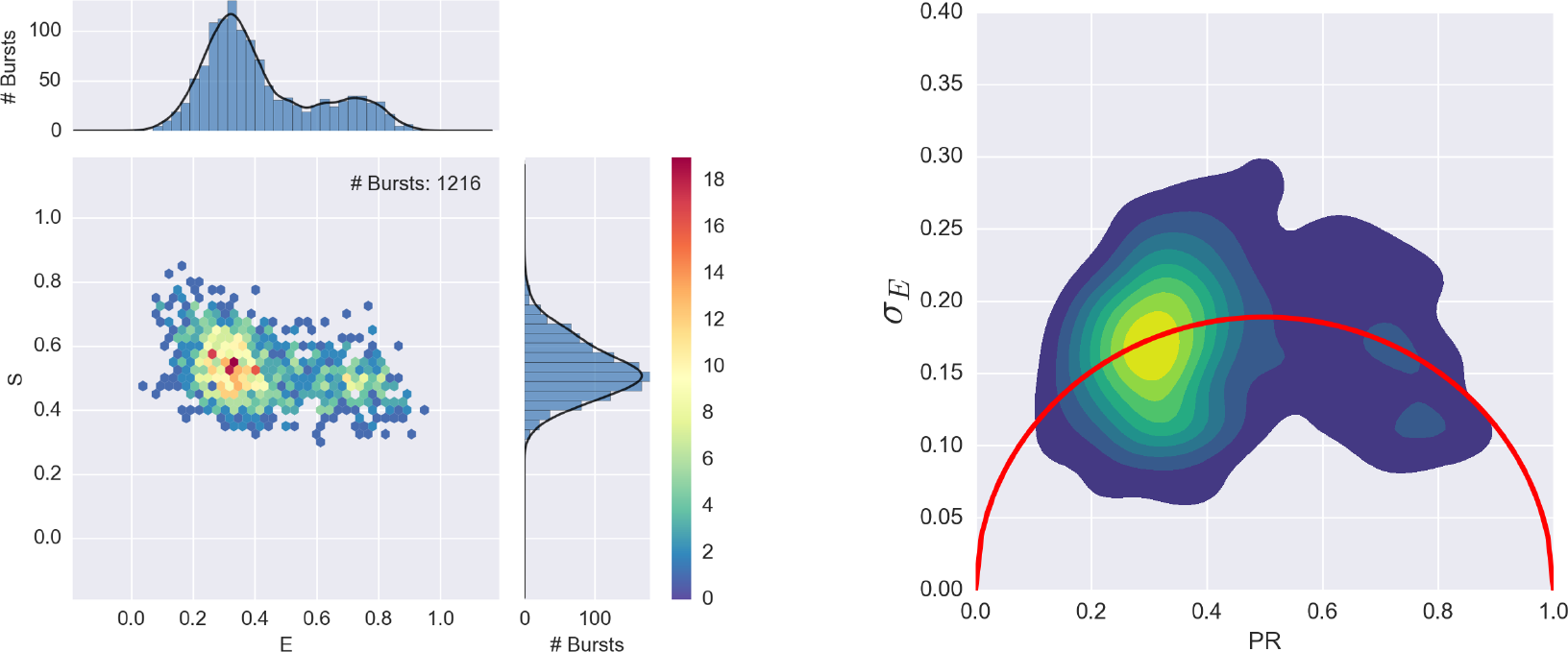
**BVA distribution for a static mixture sample**. The left panel shows the E-S histogram for a mixture of single stranded DNA (20dT) and double stranded DNA (20dT-20dA) molecules in 200 mM MgCl_2_. The right panel shows the corresponding BVA plot. Since both 20dT and 20dT-20dA are stable and have no dynamics, the BVA plots shows *s_E_* peaks lying on the static standard deviation curve (*red curve*).

**Figure 8:**
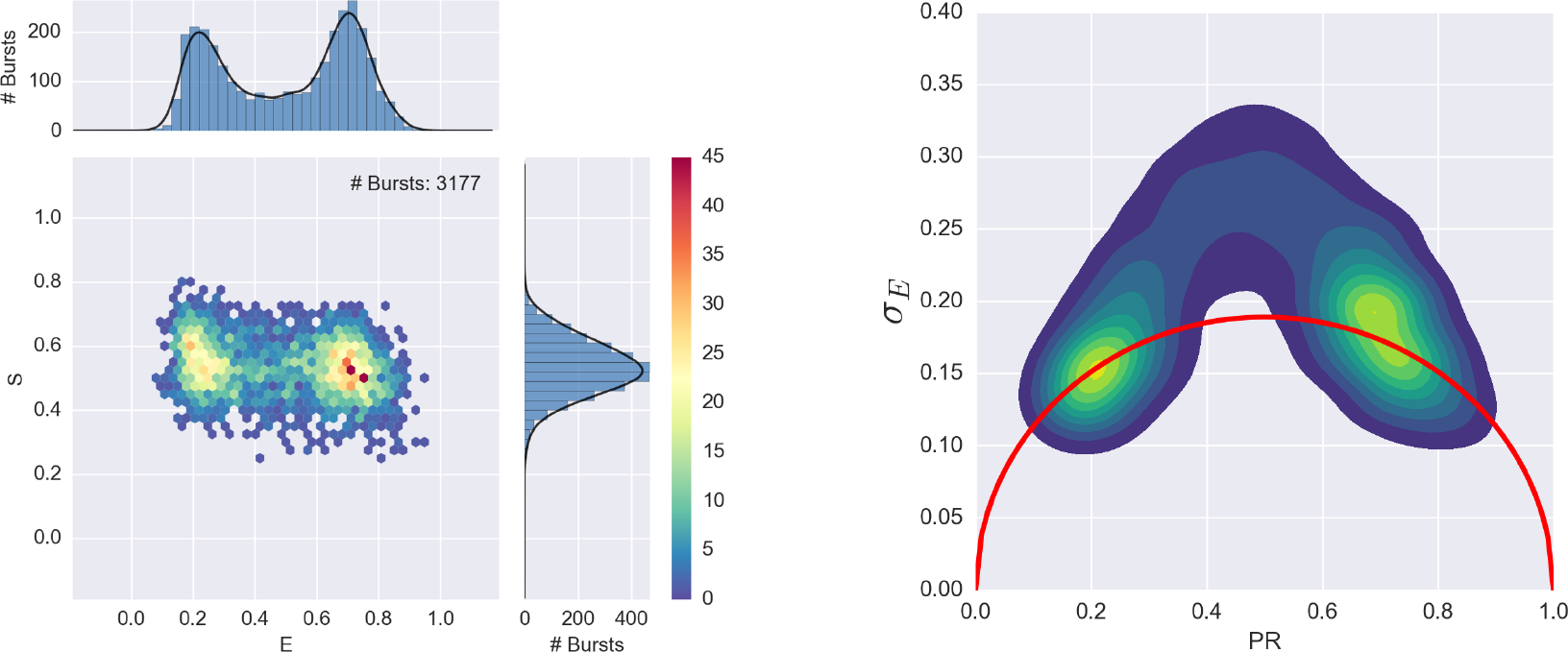
**BVA distribution for a hairpin sample undergoing dynamics**. The left panel shows the E-S histogram for a single stranded DNA sample (*A_3_*1-TA, see in [68]), designed to form a transient hairpin in 400mM NaCl. The right panel shows the corresponding BVA plot. Since the transition between hairpin and open structure causes a significant change in FRET efficiency, s_E_ lies largely above the static standard deviation curve (*red curve*).

Here, *n* is the BVA parameter defining the number of photons in each burst chunk. The outer loop iterates through bursts, while the inner loop iterates through sub-bursts. The variables `startlist` and `stoplist` are the list of start-stop indexes for all sub-bursts in current burst. In the inner loop, `A_D` and `E` contain the number of acceptor photons and FRET efficiency for the current sub-burst. Finally, for each burst, the standard deviation of E is appended to the list `E_sub_std`.

By plotting the 2D distribution of *s_E_* (i.e. `E_sub_std`) versus the average (uncorrected) E we obtain the BVA plots of figure 7 and 8.

## 6 Conclusions

FRETBursts is an open source and openly developed (see SI S2) implementation of established smFRET burst analysis methods made available to the single-molecule community. It implements several novel concepts which improve the analysis results, such as time-dependent background estimation, background-dependent burst search threshold, burst weighting and γ-corrected burst size selection. More importantly, FRETBursts provides a library of thoroughly-tested functions for timestamps and burst manipulation, making it an ideal platform for developing and comparing new analytical techniques.

We envision FRETBursts both as a state-of-the-art burst analysis software as well as a platform for development and assessment of novel algorithms. To underpin this envisioned role, FRETBursts is developed following modern software engineering practices, such as DRY principle (link) to reduce duplication and KISS principle (link) to reduce overengineering. Furthermore, to minimize the number software errors [36,69], we employ defensive programming [39] which includes code readability, unit and regression testing and continuous integration [28]. Finally, being open source, any scientist can inspect the source code, fix errors, adapt it to her own needs.

We believe that, in the single-molecule community, standard open source software implementations, such as FRETBursts, can enhance reliability and reproducibility of analysis and promote a faster adoption of novel methods, while reducing the duplication of efforts among different groups.

## Acknowledgments

We thank Dr. Eyal Nir and Dr. Toma Tomov for support in the implementation of the 2CDE method and Dr. Achillefs Kapanidis and Dr. Nicole Robb for providing experimental data for testing the BVA implementation. This work was supported by National Institutes of Health (NIH) grant R01-GM95904 and R01-GM069709. Dr. Weiss discloses equity in Nesher Technologies and intellectual property used in the research reported here. The work at UCLA was conducted in Dr. Weiss’s Laboratory.

## Supporting Information

### S1 Notebook Workflow

FRETBursts has been developed with the goal of facilitating computational reproducibility of the performed data analysis [25]. For this reason, the preferential way of using FRETBursts is by executing one of the tutorials which are in the form of Jupyter notebooks [35]. Jupyter (formerly IPython) notebooks are web-based documents which contain both code and rich text (including equations, hyperlinks, figures, etc…). FRETBursts tutorials are notebooks which can be re-executed, modified or used to process new data files with minimal modifications. The “notebook workflow” [35] not only facilitates the description of the analysis (by integrating the code in a rich document) but also greatly enhances its reproducibility by storing an execution trail that includes software versions, input files, parameters, commands and all the analysis results (text, figures, tables, etc.).

The Jupyter Notebook environment streamlines FRETBursts execution (compared to a traditional script and terminal based approach) and allows FRETBursts to be used even without prior python knowledge. The user only needs to get familiar with the notebook graphical environment, in order to be able to navigate and run the notebooks. A list of all FRETBursts notebooks can be found in the FRETBursts_notebooks repository on GitHub(link). Finally, we provide a service to run FRETBursts notebooks online, without requiring any software installation (link).

### S2 Development and Contributions

Errors are an inevitable reality in any reasonably complex software [36,69]. It is therefore critical to implement countermeasures to minimize the probability of introducing bugs and their potential impact [37,39]. In developing FRETBursts we leverage open source technlogies and follow modern software development best-practices as summarized below.

FRETBursts (and the entire python ecosystem it depends on) is open source and the source code is fully available for any scientist to study, review and modify. The open source nature of FRETBursts and of the python ecosystem, not only makes it a more transparent, reviewable platform for scientific data analysis, but also allows to leverage state-of-the-art online services such as GitHub (link) for hosting, issues tracking and code reviews, TravisCI (link) and AppVeyor (link) for continuous integration (i.e. automated test suite execution on multiple platforms after each commit) and ReadTheDocs.org for automatic documentation building and hosting. All these services would be extremely costly, if available at all, for a proprietary software or platform [70].

We highly value source code readability, a property which can reduce the number of bugs by facilitating understanding and verifying the code. For this purpose, FRETBursts code-base is well commented (with comments representing over 35% of the source code), follows the PEP8 python code style rules (link), and has docstrings in napoleon format (link).

Reference documentation is built with Sphinx (sphinx-doc.org) and all API documents are automatically generated from docstrings. On each commit, documentation is automatically built and deployed on ReadTheDocs.org.

Unit tests cover most of the core algorithms, ensuring consistency and minimizing the probability of introducing bugs. The continuous integration services, execute the full test suite on each commit on multiple platforms, immediately reporting errors. As a rule, whenever a bug is discovered, the fix also includes a new test to ensure that the same bug does not happen in the future. In addition to the unit tests, we include a regression-test notebook (link) to easily compares numerical results between two versions of FRETBursts. Additionally, the tutorials themselves are executed before each release as an additional test layer to ensure that no errors or regressions are introduced.

RETBursts is openly developed using the GitHub platform. The authors encourage users to use GitHub issues for questions, discussions and bug reports, and to submit patches through GitHub pull requests. Contributors of any level of expertise are welcome in the projects and publicly acknowledged. Contributions can be as simple as pointing out deficiencies in the documentation but can also be bug reports or corrections to the documentation or code. Users willing to implement new features are encouraged to open an Issue on GitHub and to submit a Pull Request. The open source nature of FRETBursts guarantees that contributions will become available to the entire single-molecule community.

### S3 Timestamps and Burst Data

Beyond providing prepackaged functions for established methods, FRETBursts also provides the infrastructure for exploring new analysis approaches. Users can easily get timestamps (or selection masks) for any photon stream. Core burst data (start and stop times, indexes and derived quantities for each burst) are stored in Bursts objects (link). This object provides a simple and well-tested interface (100 % unit-test coverage) to access and manipulate burst data. Bursts are created from a sequence of start/stop times and indexes, while all other fields are automatically computed. Bursts’s methods allow to recompute indexes relative to a different photon selection or recompute start and stop times relative to a new timestamps array. Additional methods perform fusion of nearby bursts or combination of two set of bursts (time intersection or union).This functionality is used for example to implement the DCBS. In conclusion, Bursts efficiently implements all the common operations performed with burst data, providing and easy-to-use interface and well tested algorithms. Leveraging Bursts methods, users can implement new types of analysis without wasting time implementing (and debugging) standard manipulation routines. Examples of working directly with timestamps, masks (i.e. photon selections) and burst data are provided in one of the FRETBursts notebooks (link). Section 5 provides a complete example on using FRETBursts to implement custom burst analysis techniques.

**Python details** Timestamps are stored in the Data attribute `ph_times_m`, which is a list or arrays, one array per excitation spot. In single-spot measurements the full timestamps array is accessed as `Data.ph_times_m`[0]. To get timestamps of arbitrary photon streams, users can call `Data.get_ph_times` (link). Photon streams are selected from the full (all-photons) timestamps array using boolean masks, which can be obtained calling `Data.get_ph_mask` (link). All burst data (e.g. start-stop times and indexes, burst duration, etc.) are stored in Bursts objects. For uniformity, the bursts start-stop indexes are always referring to the all-photons timestamps array, regardless of the photon stream used for burst search. Bursts objects internally store only start and stop times and indexes. The other Bursts attributes (duration, photon counts, etc.) are computed on-the-fly when requested (using class properties), thus minimizing the object state. Bursts support iteration with performances similar to iterating through rows of 2D row-major numpy arrays.

### S4 Plotting Data

FRETBursts uses matplotlib [47] and seaborn [71] to provide a wide range of built-in plot functions (link) for Data objects. The plot syntax is the same for both single and multi-spot measurements. The majority of plot commands are called through the wrapper function dplot, for example to plot a timetrace of the photon data, type:

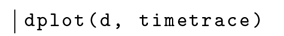

The function dplot is the generic plot function, which creates figure and handles details common to all the plotting functions (for instance, the title). d is the Data variable and timetrace is the actual plot function, which operates on a single channel.In multispot measurements, dplot creates one subplot for each spot and calls timetrace for each channel.

All built-in plot functions which can be passed to dplot are defined in the `burst_plot` module (link).

**Python details** When FRETBursts is imported, all plot functions are also imported. To facilitate finding the plot functions through auto-completion, their names start with a standard prefix indicating the plot type. The prefixes are: `timetrace` for binned timetraces of photon data, `ratetrace` for rates of photons as a function of time (non binnned), hist for functions plotting histograms and `scatter` for scatter plots. Additional plots can be easily created directly with matplotlib.

By default, in order to speed-up batch processing, FRETBursts notebooks display plots as static images using the *inline* matplotlib backend. User can switch to interactive figures inside the browser by activating the interactive backend with the command `%matplotlib` notebook. Another option is displaying figures in a new standalone window using a desktop graphical library such as QT4. In this case, the command to be used is `%matplotlib` qt.

A few plot functions, such as `timetrace` and `hist2d_alex`, have interactive features which require the QT4 backend. As an example, after switching to the QT4 backend the following command:

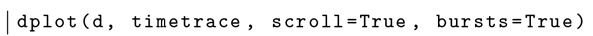
will open a new window with a timetrace plot with overlay of bursts, and an horizontal scroll-bar for quick “scrolling” throughout time. The user can click on a burst to have the corresponding burst info be printed in the notebook. Similarly, calling the hist2d_alex function with the QT4 backend allows selecting an area on the E-S histogram using the mouse.

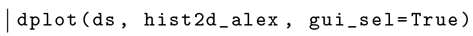

The values which identify the region are printed in the notebook and can be passed to the function select_bursts.ES to select bursts inside that region (see section *Burst Selection* in the main text).

### Plotting ALEX histograms

E-S histograms are traditionally computed using a bin size of 0.02-0.04, and cover a range slightly larger than the [0, 1] interval in which ratios of quantities not corrected for background would normally fall. FRETBursts allows plotting the square-bin 2-D E-S histogram using the plot function hist2d_alex as shown in the previous example. The histogram can be “smothed” via bicubic interpolation between bin centers (default) or plotted with raw square bins (pass the argument interpolation=’none’). Additionally, a scatter plot of E-S points is by default overlayed and it can be useful for observing the burst distribution in sparse regions. However, the different layers make this plot hard to read.

A more elegant approach for effectively representing E-S histograms with minimal clutter and high information content is using an hexbin plot (as used by the FRETBursts function `alex_jointplot`). The hexbin plot is a 2-D histograms using hexagonal bins that reduces gridding artifacts of square bins. In addition, in sparse regions, the hexbin plot naturally resembles a scatter plot (with hexagonal markers). The use of hexagonal bins for 2D distributions has been pioneered by Dan Carr in S-PLUS and then popularized by Nicholas Lewin-Koh which wrote the R language port. Later, hexbin has been implemented in the matplotlib python library (which is what FRETBursts uses). The advantages of hexagonal bins have been extensively studied and can be summarized with Nicholas Lewin-Koh words (link):

> Why hexagons? There are many reasons for using hexagons, at least over squares. Hexagons have symmetry of nearest neighbors which is lacking in square bins. Hexagons are the maximum number of sides a polygon can have for a regular tesselation of the plane, so in terms of packing a hexagon is 13% more efficient for covering the plane than squares. This property translates into better sampling efficiency at least for elliptical shapes. Lastly hexagons are visually less biased for displaying densities than other regular tesselations.

The function `alex_jointplot plots` a 3-panels plots with a central hexbin plot of E-S values and marginal E and S histograms represented in top and right panel (see figure 4 and 5 in the main text). Note that unlike other plot functions in FRETBursts, `alex_jointplot` is called directly and not through the `dplot` wrapper.

## S5 Background Estimation With Optimal Threshold

The functions used to fit the background (i.e. bg.exp_fit and other functions in bg module) provide also a goodness-of-fit estimator computed from the empirical distribution function (EDF) [72, 73]. The “distance” between the EDF and the theoretical (i.e. exponential) cumulative distribution represents and indicator of the quality of fit. Two different distance metrics can be returned by the background fitting functions. The first is the Kolgomorov-Smirnov statistics, which uses the maximum of the difference between the EDF and the theoretical distribution. The second is the Cramer von Mises statistics corresponding to the integral of the squared residuals (see the code for more details, link).

In principle, the optimal inter-photon delay threshold will minimize the error metric. This approach is implemented by the function calc_bg_brute (link) which performs a brute-force search in order to find the optimal threshold. This optimization is not necessary under typical experimental conditions, because the estimated rates normally change only a by a few per-cent compared to the heuristic threshold selection used by default.

## S6 Burst Weights

### S6.1 Theory

Freely-diffusing molecules across a Gaussian excitation volume give rise to a burst size distribution that is exponentially distributed. In a static FRET population, burst counts in the acceptor channel can be modeled as a binomial random variable (RV) with success probability equal to the population PR and number of trials equal to the burst size *n_d_* + *n_a_*. Similarly, the PR of each burst *E_i_* (*i* being the burst index) is simply a binomial divided by the number of trials, with variance reported in eq. 4.

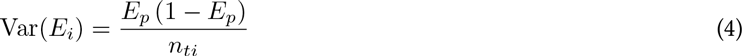

Bursts with higher counts, provide more accurate estimations of the population PR, since their PR variance is smaller (eq. 4). Therefore, in estimating the population PR we need to “focus” on bigger bursts. Traditionally, this is accomplished by merely discarding bursts below a size-threshold. In the following paragraphs we demonstrate how, by proper weighting bursts, is possible to obtains optimal estimates of PR which takes into account the information of the entire burst population.

According to the Cramer-Rao lower bound (eq. 5), the Fisher information *I*(θ) sets a lower bound on the variance of any statistics 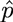 of a RV θ.

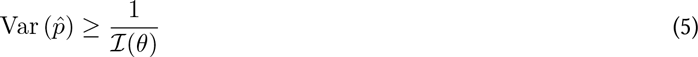

When the statistics 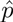 is an unbiased estimator of a distribution parameter and the equality holds in eq. 5, the estimator is a minimum-variance unbiased (MVUB) estimator and it is said to be efficient (meaning that it does an optimal use the information contained in the sample to estimate the parameter).

A population of *N* bursts can be modeled by a set of *N* binomial variables with same success probability *E_p_* and varying number of successes equal to the burst size. An estimator for *E_p_* can be constructed noticing that the sum of binomial RV with same success probability is still a binomial (with number of trials equal to the sum of the number of trials). Taking the sum of acceptor counts over all bursts divided by the total number of photons as in eq. 6, we obtain an estimator *Ê* of the proportion of successes

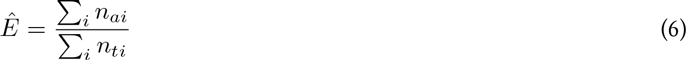

The variance of *Ê* (eq. 7) is equal to the inverse of the Fisher information I(*Ê*) and therefore *Ê* is a MVUB estimator for *E_p_*.

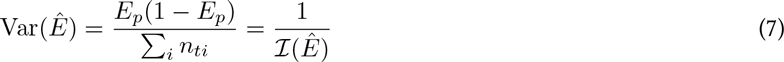

We can finally verify that *Ê* is equal to the weighted average of the bursts PR *E_i_* (eq. 9), with weights proportional to the burst size (eq. 8).

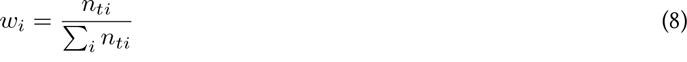

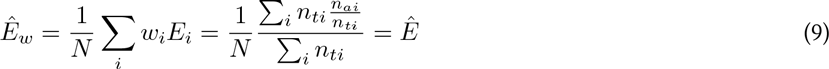

Since *Ê* is the MVUB estimator, any other estimator of *E_p_* (in particular the unweighted mean of *E*_*i*_) will have a larger variance.

We can extend these consideration of optimal weights for the PR estimator to the FRET distribution plot (histograms or KDEs). Building an unweighted histogram (and fitting the peak) is analogous to estimating the *E_p_* with an unweighted average. Conversely, building the FRET histogram using the burst size as weights is equivalent to using the MVUB estimator for *E_p_*.

### S6.2 Weighted FRET Estimator

Here we report a simple verification of the results of previous section, namely that a weighted mean of *E_i_* is the estimator with minimal variance of *E_p_*. For this purpose, we generated a static FRET population of 100 bursts by simply extracting burst-sizes from an exponential distribution (*λ* = 10) and acceptor counts from a binomial distribution (*E_p_* = 0.2). By repeatedly fitting the population parameter *E*_*p*_ using a size-weighted and unweighted average, we verified that the former has systematically lower variance of the latter as predicted by the theory (in the current example the unweighted estimator has 28.6% higher variance). Note that this result holds for any arbitrary distribution of burst sizes. The full simulation including exponential and gamma-distributed burst sizes is reported in the accompanying Jupyter notebook (link).

### S6.3 Weighted FRET Histogram

The effect of weighting the FRET histogram is here illustrated with a simulation of a mixture of two static FRET populations and then with experimental data.

We performed a realistic simulation of a static mixture of two FRET populations starting from 3-D Brownian motion diffusion of *N* particles excited by a numerically computed (non-Gaussian) PSF. Input parameters of the simulation include diffusion coefficient, particle brightness, the two FRET efficiencies, as well as detectors DCR. The simulation is performed with the open source software PyBroMo [41] which creates smFRET data files (i.e. timestamps and detectors arrays) in Photon-HDF5 format [40]. The simulated data file is processed with FRETBursts performing burst search, and only a minimal burst size selection of with threshold of 10 photons. The resulting weighted and unweighted FRET histograms are reported in figure S1. We notice that the use of the weights results in better definition of FRET peaks.

As a final comparison, we report the weighted and unweighted FRET histogram of an experimental FRET population from measurement of a di-labeled dsDNA sample. Figure S2 show a comparison of a FRET histogram obtained from the same burst with and without weights. The burst selection is obtained applying a burst size threshold of 10 counts (after background correction), in order to filter the extreme low-end of the burst size distribution.

The use of size-weighted FRET histograms is a simple way to obtain a representation of FRET distribution that maintains high power of resolving FRET peaks while including the full burst population and thus reducing statistical noise.

As a final remark, note that when increasing the size-threshold for burst selection the difference between weighted and unweighted FRET histograms tends to zero because the relative difference in burst weights in the selected burst becomes smaller (i.e. tends to 1, meaning equal weights).

**Figure S1:**
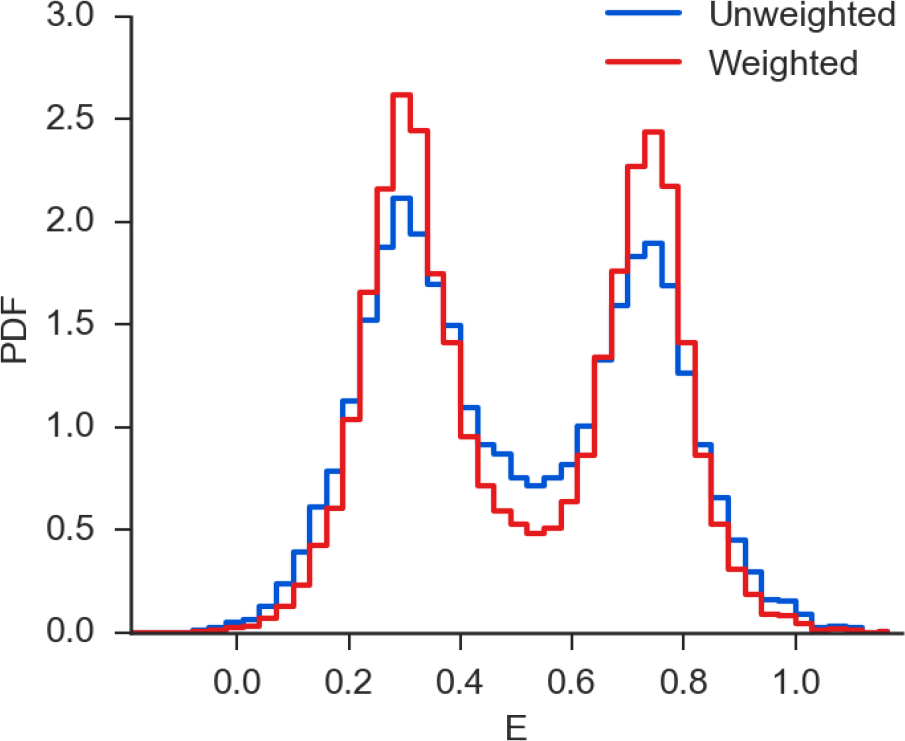
Comparison of unweighted and size-weighted FRET histograms for a simulated mixtures of static FRET populations. In both cases bursts are selected with a size threshold of 10 photons (after background correction)

**Figure S2:**
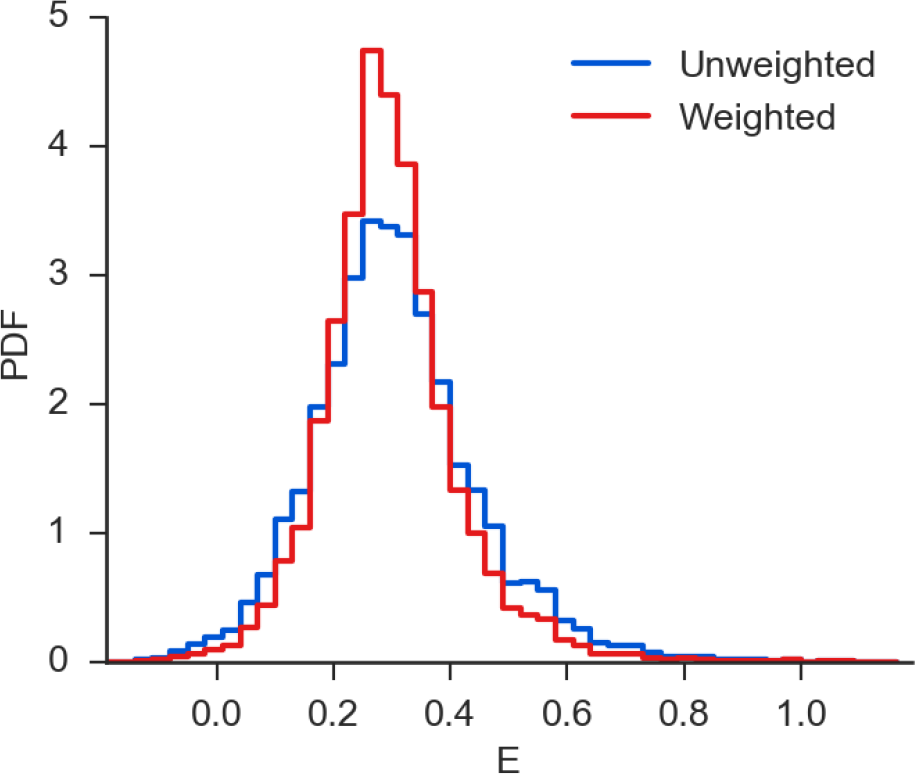
Comparison of unweighted and size-weighted FRET histograms for a smFRET measurement of a static FRET sample (di-labeled dsDNA). In both cases bursts are selected with a size threshold of 10 photons (after background correction)

## References

[1] S. Weiss. Fluorescence Spectroscopy of Single Biomolecules. Science, 283(5408):1676–1683, mar 1999. 10.1126/science.283.5408.1676.

[2] JohannesHohlbein, TimothyD. Craggs, and Thor-benCordes. Alternating-laser excitation: single molecule FRET and beyond. Chemical Society Reviews, 43(4):1156–1171, January 2014. 10.1039/C3CS60233H.

[3] E Lerner, T Orevi, E Ben Ishay, D Amir, and E Haas. Kinetics of fast changing intramolecular distance distributions obtained by combined analysis of FRET efficiency kinetics and time-resolved FRET equilibrium measurements. Biophysical Journal, 106(3):667–76, feb 2014. 10.1016/j.bpj.2013.11.4500.

[4] Gil Rahamim, Marina Chemerovski-Glikman, Shai Rahimipour, Dan Amir, and ElishaHaas. Resolution of Two Sub-Populations of Conformers and Their Individual Dynamics by Time Resolved Ensemble Level FRET Measurements. PloS ONE, 10(12):e0143732, jan 2015. 10.1371/journal.pone.0143732.

[5] Paul R. Selvin. The renaissance of fluorescence resonance energy transfer. Nature Structural & Molecular Biology, 7(9):730–734, September 2000. 10.1038/78948.

[6] RahulRoy, SungchulHohng, and TaekjipHa. A practical guide to single-molecule FRET. Nature Methods, 5(6):507–516, jun 2008. 10.1038/nmeth.1208.

[7] Benjamin Schuler and William A Eaton. Protein folding studied by single-molecule FRET. Current Opinion in Structural Biology, 18(1):16–26, feb 2008. 10.1016/j.sbi.2007.12.003.

[8] EvangelosSisamakis, AlessandroValeri, StanislavKalinin, Paul J Rothwell, and C. A. M. Seidel. Accurate Single-Molecule FRET Studies Using Multiparameter Fluorescence Detection. In Methods in Enzymology, volume 475, pages 455–514. Elsevier Inc., 1 edition, January 2010. 10.1016/s0076-6879(10)75018-7.

[9] Gilad Haran. How when and why proteins collapse: the relation to folding. Current Opinion in Structural Biology, 22(1):14–20, feb 2012. 10.1016/j.sbi.2011.10.005.

[10] C. Eggeling, J. R. Fries, L. Brand, R. Gunther, and C. A. M. Seidel. Monitoring conformational dynamics of a single molecule by selective fluorescence spectroscopy. Proceedings of the National Academy of Sciences USA, 95(4):1556–1561, feb 1998. 10.1073/pnas.95.4.1556.

[11] Maxime Dahan, Ashok A. Deniz, TaekjipHa, Daniel S. Chemla, Peter G. Schultz, and Shimon Weiss. Ratiometric measurement and identification of single diffusing molecules. Chemical Physics, 247(1):85–106, aug 1999. 10.1016/s0301-0104(99)00132-9.

[12] Joachim R. Fries, Leif Brand, ChristianEggeling, Malte Kollner, and Claus A. M. Seidel. Qantitative Identification of Different Single Molecules by Selective Time— Resolved Confocal Fluorescence Spectroscopy. The Journal of Physical Chemistry A, 102(33):6601–6613, August 1998. 10.1021/jp980965t.

[13] C. Eggeling, S. Berger, L. Brand, J.R. Fries, J. Schaffer, A. Volkmer, and C.A.M. Seidel. Data registration and selective single–molecule analysis using multi-parameter fluorescence detection. Journal of Biotechnology, 86(3):163–180, apr 2001. 10.1016/s0168-1656(00)00412-0.

[14] Kai Zhang and Haw Yang. Photon–by–Photon Determination of Emission Bursts from Diffusing Single Chromophores. The Journal of Physical Chemistry B, 109(46):21930–21937, November 2005. 10.1021/jp0546047.

[15] Irina Gopich and AttilaSzabo. Theory of photon statistics in single–molecule Forster resonance energy transfer. The Journal of Chemical Physics, 122(1):014707, 2005. 10.1063/1.1812746.

[16] Nam KiLee, Achillefs N. Kapanidis, You Wang, Xavier Michalet, Jayanta Mukhopadhyay, Richard H. Ebright, and Shimon Weiss. Accurate FRET Measurements within Single Diffusing Biomolecules Using Alternating–Laser Excitation. Biophysical Journal, 88(4):2939–2953, apr 2005. 10.1529/biophysj.104.054114.

[17] Eyal Nir, Xavier Michalet, Kambiz M. Hamadani, Ted A. Laurence, Daniel Neuhauser, Yevgeniy Kovchegov, and Shimon Weiss. Shot-Noise Limited Single-Molecule FRET Histograms– Comparison between Theory and Experiments. The Journal of Physical Chemistry B, 110(44):22103–22124, nov 2006. 10.1021/jp063483n.

[18] Matthew Antonik, Suren Felekyan, Alexander Gaiduk, and Claus A M Seidel. Separating structural heterogeneities from stochastic variations in fluorescence resonance energy transfer distributions via photon distribution analysis. Journal of Physical Chemistry B, 110(13):6970–6978, apr 2006. 10.1021/jp057257+.

[19] Irina V. Gopich and Attila Szabo. Single-Molecule FRET with Diffusion and Conformational Dynamics. The Journal of Physical Chemistry B, 111(44):12925–12932, nov 2007. 10.1021/jp075255e.

[20] Irina V. Gopich. Concentration Effects in “SingleMolecule” Spectroscopy. The Journal ofPhysical Chemistry B, 112(19):6214–6220, may 2008. 10.1021/jp0764182.

[21] Brian A. Camley, Frank L. H. Brown, and Everett A. Lip– man. Förster transfer outside the weak-excitation limit. The Journal of Chemical Physics, 131(10):104509, 2009. 10.1063/1.3230974.

[22] Yusdi Santoso, Joseph P. Torella, and Achillefs N. Kapanidis. Characterizing Single–Molecule FRET Dynamics with Probability Distribution Analysis. ChemPhysChem, 11(10):2209–2219, jun 2010. 10.1002/cphc.201000129.

[23] Joseph P. Torella, Seamus J. Holden, Yusdi Santoso, Johannes Hohlbein, and Achillefs N. Kapanidis. Identifying Molecular Dynamics in Single–Molecule FRET Experiments with Burst Variance Analysis. Biophysical Journal, 100(6):1568–1577, mar 2011. 10.1016/j.bpj.2011.01.066.

[24] Toma E. Tomov, Roman Tsukanov, Rula Masoud, Miran Liber, Noa Plavner, and Eyal Nir. Disentangling Subpopulations in Single–Molecule FRET and ALEX Experiments with Photon Distribution Analysis. Biophysical Journal, 102(5):1163–1173, mar 2012. 10.1016/j.bpj.2011.11.4025.

[25] Jonathan B. Buckheit and David L. Donoho. WaveLab and Reproducible Research. In Wavelets and Statistics, volume 103, pages 55–81. Springer Science + Business Media, 1995. 10.1007/978-1-4612-2544-7_5.

[26] Darrel C. Ince, Leslie Hatton, and John Graham– Cumming. The case for open computer programs. Nature, 482(7386):485–488, feb 2012. 10.1038/nature10836.

[27] Mauno Vihinen. No more hidden solutions in bioinformatics. Nature, 521(7552):261–261, may 2015. 10.1038/521261a.

[28] Stephen Eglen, Ben Marwick, Yaroslav Halchenko, Michael Hanke, Shoaib Sufi, Padraig Gleeson, R. Angus Silver, Andrew Davison, Linda Lanyon, Mathew Abrams, Thomas Wachtler, David J Willshaw, Christophe Pouzat, and Jean–Baptiste Poline. Towards standard practices for sharing computer code and programs in neuroscience. bioRxiv, mar 2016. 10.1101/045104.

[29] Sean A. McKinney, Chirlmin Joo, and Taekjip Ha. Analysis of Single–Molecule FRET Trajectories Using Hidden Markov Modeling. Biophysical Journal, 91(5):1941–1951, sep 2006. 10.1529/biophysj.106.082487.

[30] Jonathan E. Bronson, Jingyi Fei, Jake M. Hofman, Ruben L. Gonzalez, and Chris H. Wiggins. Learning Rates and States from Biophysical Time Series: A Bayesian Approach to Model Selection and SingleMolecule FRET Data. Biophysical Journal, 97(12):3196–3205, dec 2009. 10.1016/j.bpj.2009.09.031.

[31] Max Greenfeld, Dmitri S. Pavlichin, Hideo Mabuchi, and Daniel Herschlag. Single Molecule Analysis Research Tool (SMART): An Integrated Approach for Analyzing Single Molecule Data. PLoS ONE, 7(2):e30024, feb 2012. 10.1371/journal.pone.0030024.

[32] Sebastian L. B. Konig, Melodie Hadzic, Erica Fiorini, Richard Borner, Danny Kowerko, Wolf U. Blancken– horn, and Roland K. O. Sigel. B0BA FRET: Bootstrap– Based Analysis of Single–Molecule FRET Data. PLoS ONE, 8(12):e84157, dec 2013. 10.1371/journal.pone.0084157.

[33] Jan–Willem van de Meent, Jonathan E. Bronson, Chris H. Wiggins, and Ruben L. Gonzalez. Empirical Bayes Methods Enable Advanced Population–Level Analyses of Single-Molecule FRET Experiments. Biophysical Journal, 106(6):1327–1337, mar 2014. 10.1016/j.bpj.2013.12.055.

[34] Rebecca R. Murphy, Sophie E. Jackson, and David Klen– erman. pyFRET: A Python Library for Single Molecule Fluorescence Data Analysis. ArXiv, dec 2014. arXiv:1412.6402.

[35] Helen Shen. Interactive notebooks: Sharing the code. Nature, 515(7525):151–152, nov 2014. 10.1038/515151a.

[36] David A. W. Soergel. Rampant software errors may undermine scientific results. F1000Research, jul 2015. 10.12688/f1000research.5930.2.

[37] Greg Wilson, D. A. Aruliah, C. Titus Brown, Neil P. Chue Hong, Matt Davis, Richard T. Guy, Steven H. D. Haddock, Kathryn D. Huff, Ian M. Mitchell, Mark D. Plumbley, Ben Waugh, Ethan P. White, and Paul Wilson. Best Practices for Scientific Computing. PLoS Biology, 12(1):e1001745, jan 2014. 10.1371/journal.pbio.1001745.

[38] John D. Blischak, Emily R. Davenport, and Greg Wilson. A Quick Introduction to Version Control with Git and GitHub. PLOS Computational Biology, 12(1):e1004668, jan 2016. 10.1371/journal.pcbi.1004668.

[39] Andreas Prlic and James B. Procter. Ten Simple Rules for the Open Development of Scientific Software. PLoS Computational Biology, 8(12):e1002802, dec 2012. 10.1371/journal.pcbi.1002802.

[40] Antonino Ingargiola, Ted Laurence, Robert Boutelle, Shimon Weiss, and Xavier Michalet. Photon–HDF5: An 0pen File Format for Timestamp-Based SingleMolecule Fluorescence Experiments. Biophysical Journal, 110(1):26–33, jan 2016. 10.1016/j.bpj.2015.11.013.

[41] Antonino Ingargiola, Ted Laurence, Robert Boutelle, Shimon Weiss, and Xavier Michalet. Photon– HDF5: open data format and computational tools for timestamp–based single–molecule experiments. volume 971405. SPIE, 2016. 10.1117/12.2212085.

[42] Antonino Ingargiola, Francesco Panzeri, Niusha Sarkhosh, Angelo Gulinatti, Ivan Rech, Massimo Ghioni, Shimon Weiss, and Xavier Michalet. 8–spot smfret analysis using two 8–pixel spad arrays. volume 8590, pages 85900E–85900E-11. SPIE, 2013. 10.1117/12.2003704.

[43] Achillefs N. Kapanidis, Ted A. Laurence, Nam Ki Lee, Emmanuel Margeat, Xiangxu Kong, and Shimon Weiss. Alternating–Laser Excitation of Single Molecules. Accounts of Chemical Research, 38(7):523–533, jul 2005. 10.1021/ar0401348.

[44] Ted A. Laurence, Xiangxu Kong, Marcus Jäger, and Shimon Weiss. Probing structural heterogeneities and fluctuations of nucleic acids and denatured proteins. Proceedings of the National Academy of Sciences USA, 102(48):17348–17353, 2005. 10.1073/pnas.0508584102.

[45] Barbara K. Muller, Evgeny Zaychikov, Christoph Braauchle, and Don C. Lamb. Pulsed Interleaved Excitation. Biophysical Journal, 89(5):3508–3522, nov 2005. 10.1529/biophysj.105.064766.

[46] X. Michalet, R. A. Colyer, G. Scalia, A. Ingargiola, R. Lin, J. E. Millaud, S. Weiss, O. H. W. Siegmund, A. S. Tremsin, J. V. Vallerga, A. Cheng, M. Levi, D. Aha– roni, K. Arisaka, F. Villa, F. Guerrieri, F. Panzeri, I. Rech, A. Gulinatti, F. Zappa, M. Ghioni, and S. Cova. Development of new photon–counting detectors for single–molecule fluorescence microscopy. Philosophical Transactions of the Royal Society B: Biological Sciences, 368(1611):20120035–20120035, dec 2012. 10.1098/rstb.2012.0035.

[47] Michael Droettboom, John Hunter, Thomas A Caswell, Eric Firing, Jens Hedegaard Nielsen, Phil Elson, Benjamin Root, Darren Dale, Jae–JoonLee, Jouni K. Seppaanen, Damon McDougall, Andrew Straw, Ryan May, Nelle Varoquaux, TonyS Yu, EricMa, CharlieMoad, Steven Silvester, ChristophGohlke, Peter Wurtz, Thomas Hisch, FedericoAriza, Cimarron, Ian Thomas, James Evans, Paul Ivanov, Jeff Whitaker, Paul Hobson, mdehoon, and Matt Giuca. matplotlib: matplotlib v1.5.1, jan 2016. 10.5281/zenodo.44579.

[48] L. Edman, U. Mets, and R. Rigler. Conformational transitions monitored for single molecules in solution. Proceedings of the National Academy of Sciences USA, 93(13):6710–6715, jun 1996. 10.1073/pnas.93.13.6710.

[49] A. A. Deniz, M.Dahan, J. R. Grunwell, T. Ha, A. E. Faulhaber, D. S. Chemla, S. Weiss, and P. G. Schultz. Single–pair fluorescence resonance energy transfer on freely diffusing molecules: Observation of Forster distance dependence and subpopulations. Proceedings of the National Academy of Sciences USA, 96(7):3670–3675, mar 1999. 10.1073/pnas.96.7.3670.

[50] A. N. Kapanidis, N. K. Lee, T. A. Laurence, S. Doose, E. Margeat, and S. Weiss. Fluorescence–aided molecule sorting: Analysis of structure and interactions by alternating–laser excitation of single molecules. Proceedings of the National Academy of Sciences USA, 101(24):8936–8941, jun 2004. 10.1073/pnas.0401690101.

[51] Matt Newville, Andrew Nelson, Till Stensitzki, An– tonino Ingargiola, Dan Allan, Michal, YoavRam, MerlinSmiles, Li Li, Glenn, ChristophDeil, Gustavo Pasquevich, Stuermer, Tim Spillane, stonebig, Per A. Brodtkorb, Nicholas Earl, Anthony Almarza, Ben Gamari, and Kostis Anagnostopoulos. lmfit–py: release 0.9.3, April 2016. 10.5281/zenodo.49428.

[52] H. Akaike. A new look at the statistical model identification. IEEE Transactions on Automatic Control, 19(6):716–723, December 1974. 10.1109/TAC.1974.1100705.

[53] Gideon Schwarz. Estimating the Dimension of a Model. The Annals of Statistics, 6(2):461–464, March 1978. 10.1214/aos/1176344136.

[54] Stanislav Kalinin, Suren Felekyan, Matthew Antonik, and Claus A. M. Seidel. Probability Distribution Analysis of Single–Molecule Fluorescence Anisotropy and Resonance Energy Transfer. The Journal of Physical Chemistry B, 111(34):10253–10262, August 2007. 10.1021/jp072293p.

[55] Irina V. Gopich and Attila Szabo. FRET Efficiency Distributions ofMultistate Single Molecules. The Journal of Physical Chemistry B, 114(46):15221–15226, November 2010. 10.1021/jp105359z.

[56] Yusdi Santoso and Achillefs N Kapanidis. Probing biomolecular structures and dynamics of single molecules using in–gel alternating–laser excitation. Analytical chemistry, 81(23):9561–70, December 2009. 10.1021/ac901423e.

[57] Stanislav Kalinin, Evangelos Sisamakis, Steven W. Ma– gennis, Suren Felekyan, and Claus A. M. Seidel. On the origin of broadening of single–molecule fret efficiency distributions beyond shot noise limits. The Journal of Physical Chemistry B, 114(18):6197–6206, 2010. 10.1021/jp100025v.

[58] Irina V. Gopich and Attila Szabo. Decoding the pattern of photon colors in single–molecule fret. The Journal of Physical Chemistry B, 113(31):10965–10973, 2009. 10.1021/jp903671p.

[59] Irina V Gopich and Attila Szabo. Theory of SingleMolecule FRET Efficiency Histograms. In Tamiki Ko– matsuzaki, Masaru Kawakami, Satoshi Takahashi, Haw Yang, and Robert J. Silbey, editors, Single-Molecule Biophysics: Experiment and Theory, Volume 146, volume *146* of Advances in Chemical Physics, pages 245–297. John Wiley & Sons, Inc., Hoboken, NJ, USA, November 2011. 10.1002/9781118131374.ch10.

[60] ArminHoffmann, Daniel Nettels, JenniferClark, AlessandroBorgia, Sheena E Radford, JaneClarke, and Benjamin Schuler. Qantifying heterogeneity and conformational dynamics from single molecule FRET of diffusing molecules: recurrence analysis ofsingle particles (RASP). Physical Chemistry Chemical Physics, 13(5), February 2011. 10.1039/c0cp01911a.

[61] Ted A. Laurence, YoungeunKwon, EricYin, Christopher W. Hollars, Julio A. Camarero, and DanielBarsky. Correlation Spectroscopy of Minor Fluorescent Species: Signal Purification and Distribution Analysis. Biophysical Journal, 92(6):2184–2198, March 2007. 10.1529/biophysj.106.093591.

[62] TedmanTorres and Marcia Levitus. Measuring Conformational Dynamics: A New FCS-FRET Approach. The Journal of Physical Chemistry B, 111(25):7392–7400, June 2007. 10.1021/jp070659s.

[63] DanielNettels, ArminHoffmann, and BenjaminSchuler. Unfolded Protein and Peptide Dynamics Investigated with Single-Molecule FRET and Correlation Spectroscopy from Picoseconds to Seconds. The Journal of Physical Chemistry B, 112(19):6137–6146, May 2008. 10.1021/jp076971j.

[64] Matthew S. DeVore, Stephen F. Gull, and Carey K.Johnson. Classic Maximum Entropy Recovery of the Average Joint Distribution of Apparent FRET Efficiency and Fluorescence Photons for Single-Molecule Burst Measurements. The Journal of Physical Chem-istryB, 116(13):4006–4015, April 2012. doi:10.1021/jp209861u.

[65] Rebecca R.Murphy, George Danezis, Mathew H. Horrocks, Sophie E. Jackson, and DavidKlenerman. Bayesian Inference of Accurate Population Sizes and FRET Efflciencies from Single Diffusing Biomolecules. Analytical Chemistry, 86(17):8603–8612, September 2014. 0.1021/ac501188r.

[66] S. C. Kou, X. Sunney Xie, and Jun S. Liu. Bayesian analysis of single-molecule experimental data. Journal of the Royal Statistical Society: Series C (Applied Statistics), 54(3):469–506, June 2005. 10.1111/j.1467-9876.2005.00509.x.

[66] Nicole C. Robb, ThorbenCordes, Ling Chin Hwang, KristoferGryte, DiegoDuchi, Timothy D. Craggs, YusdiSantoso, ShimonWeiss, Richard H. Ebright, and Achillefs N. Kapanidis. The Transcription Bubble of the RNA Polymerase-Promoter Open Complex Exhibits Conformational Heterogeneity and Millisecond-Scale Dynamics: Implications for Transcription Start-Site Selection. Journal of Molecular Biology, 425(5):875–885, mar 2013. 10.1016/j.jmb.2012.12.015.

[68] Roman Tsukanov, Toma E. Tomov, RulaMasoud, HagaiDrory, NoaPlavner, Miran Liber, and Eyal Nir. Detailed Study of DNA Hairpin Dynamics Using SingleMolecule Fluorescence Assisted by DNA Origami. The Journal of Physical Chemistry B, 117(40):11932–11942, oct 2013. doi:10.1021/jp4059214.

[69] ZeeyaMerali. Computational science: …Error. Nature, 467(7317):775–777, oct 2010. 10.1038/467775a.

[69] JeremyFreeman. Open source tools for large-scale neuroscience. Current Opinion in Neurobiology, 32:156–163, jun 2015. 10.1016/j.conb.2015.04.002.

[70] MichaelWaskom, OlgaBotvinnik, drewokane, Paul Hobson, Yaroslav Halchenko, SauliusLukauskas, Jordi Warmenhoven, John B. Cole, StephanHoyer, JakeVan-derplas, gkunter, Santi Villalba, EricQintero, MarcelMartin, AlistairMiles, Kyle Meyer, TomAugspurger, Tal Yarkoni, PeteBachant, ConstantineEvans, Clark Fitzgerald, TamasNagy, ErikZiegler, TobiasMegies, DanielWehner, SamuelSt-Jean, Luis Pedro Coelho, GregoryHitz, AntonyLee, and LucRocher. seaborn: v0.7.0, January 2016. 10.5281/zenodo.45133.

[72] M. A. Stephens. EDF Statistics for Goodness of Fit and Some Comparisons. Journal of the American Statistical Association, 69(347):730, sep 1974. 10.2307/2286009..

[73] William C. Parr and William R. Schucany. Minimum Distance and Robust Estimation. Journal of the American Statistical Association, 75(371):616, sep 1980. 10.2307/2287658.

